# Key unprotected areas for carnivore conservation in Mexico

**DOI:** 10.1101/2024.02.10.579669

**Authors:** Germar Gonzalez, Nyeema C. Harris

## Abstract

Area-based conservation remains a pivotal component of global wildlife protection efforts. Mexico hosts a diverse array of area-based approaches, encompassing protected areas (PAs) and other effective area-based conservation measures (OECMs) such as voluntary conservation areas (VCAs) and wildlife management units (UMAs). Indigenous Territories (ITs) also provide heightened conservation potential through traditional ecological knowledge systems. These conservation spaces exhibit significant variations in community involvement, national coverage, and governance. Here, we evaluate the conservation potential of these land management types for 29 terrestrial carnivores, focusing on spatial co-occurrence. We determine areas in Mexico lacking area-based protection and calculate carnivore richness per land management type. Analyzing overlap between land management types and carnivore ranges, we employ heat maps to visualize overlap occurrence and key unprotected areas. We found that across all carnivore ranges, 87% of the area remains unprotected under designated protection areas (PAs & VCAs), decreasing to 65.2% after including non-designated types (ITs and UMAs). We identified several key gaps in the protection estate for Mexican carnivores, most notably on the eastern Mexican coast in the state of Veracruz. Our findings corroborate the importance of including Indigenous Peoples and Local Communities in conservation efforts, emphasizing their contributions to the stewardship of Mexican ecosystems. As the global protected area estate continues to expand under the post-2020 global biodiversity targets, and the importance of a well-designed and diverse portfolio of practices for conservation is recognized, the need for collective action, increased collaboration and inclusivity, and effective communication amongst stakeholders becomes necessary for carnivore conservation.

## Introduction

Area-based conservation measures such as protected areas (PAs) are at the forefront of global environmental efforts and are essential tools for combating biodiversity decline and habitat loss (Gray et al., 2016; Carroll et al., 2022). For the protection of threatened fauna in particular, PAs have become the status quo, as they help maintain prey availability, preserve habitat connectivity, and limit hunting of wildlife (Yackulic et al., 2011; Ripple et al., 2014; Chen et al., 2022). International initiatives petitioning to increase the conservation estate through global PA expansions have gained considerable momentum (eg. 30 by 30, Half-Earth), encouraged by recent studies that have located global priority biodiversity hotspots (Dinerstein et al., 2019). However, the effectiveness of PAs to mitigate anthropogenic threats and meet conservation goals in the face of growing human pressures remains under scrutiny (Brooks et al., 2004; Geldmann et al., 2019; Gurney et al., 2023). Demands to support the needs of growing human populations inherently cause conflict of space with this agenda, raising a myriad of social justice, equity, and economic concerns as conservation targets are pursued (Bennet et al., 2019; Harris et al. 2023), especially since PA expansion hotspots are clustered in the tropics throughout the Global South (Dinerstein et al., 2020). As a result, there has been a gradual shift in the conservation literature that has interrogated the traditional PA approach and recognized the potential of multiple-use, non-governmental, and non-designated conservation areas to safeguard biodiversity (Palomo et al., 2014; Butchart et al., 2014; Jonas et al., 2021). This has culminated in the conceptualization of other effective area-based conservation measures (OECMs) to complement the global PA network (Maxwell et al., 2020; Adams et al., 2023). Studies comparing traditional PA conservation potential with Indigenous Territories (ITs) or OECMs, which have community-based, bottom-up management strategies, reveal comparable outcomes in conservation performance (e.g., Elleason et al., 2021; Vimal et al., 2021). Yet, even with the growing empirical and theoretical support for OECMs in the past decades, only recently has these efforts been official recognized and now considered contributors to international biodiversity targets for PA expansion, as explicitly stated under Global Target 3 of the Kunming-Montreal Global Biodiversity Framework by the Convention on Biological Diversity (CBD) in 2022 (CBD, 2022).

Several case-studies have highlighted the positive conservation outcomes of OECMs, ITs, and other types of community conservation areas in Latin America for threatened wildlife (O’Bryan et al., 2021; Boron et al., 2022; Devlin et al., 2023). In Mexico, carnivores are a major focus of current conservation strategies because many are keystone species and contribute to ecosystem functioning (Ripple et al., 2014), hold high cultural and economic value for Indigenous Peoples and Local Communities (IPLCs) (Garcia del Valle et al., 2015; Avila-Najera et al., 2018), and are experiencing rapid population declines (Lara Diaz et al., 2023). In light of expanding human populations and increased human-wildlife interactions, conflicts are intensifying throughout Mexico’s conservation estate, necessitating more efforts to promote coexistence between humans and carnivores (Anaya-Zamora et al., 2017; Flores-Armillas et al., 2020). Assessing PA potential and effectiveness from a socio-ecological perspective is therefore essential for creating resilient and long-lasting PAs and OECMs that will complement local needs and contribute to carnivore conservation (Cumming et al., 2015; Dudley et al., 2018; Ndayizeye et al., 2020).

In a Latin American context, the inclusion of diverse area-based conservation efforts that work with IPLCs helps combat the legacies of environmental colonialism and coupled socio - ecological issues that have contributed to inequalities (Galafassi, 2010; Martinez-Alier et al., 2016). Mexico has one of the highest species diversity in Latin America, over 60 different Indigenous groups, and a mosaic of diverse area-based conservation approaches (Table 1). These diverse approaches differ in conservation designation, governance type, national area coverage, and in protection effort. Federally managed PAs are present at municipal, state, and federal levels, and there exists a range of subcategories within the PA estate, like biosphere reserves. There are also OECMs found outside of the PA estate like voluntary conservation areas (VCAs) and wildlife management units (UMAs) that employ a collaborative governance framework. ITs are also essential contributors for biodiversity conservation, and while conservation efforts may not be an intended purpose in all ITs, traditional ecological knowledge systems and their location in biodiversity hotspots often result in positive outcomes for wildlife conservation (Garnett et al., 2018; Reyes-Garcia & Benyei, 2019). These land management types often overlap and co-occur, creating a complex, multi-governance landscape across Mexico. The inclusion of non-designated land management types like ITs and UMAs as a part of the official Mexican protected area estate provide an opportunity to meet international biodiversity targets set by Target 3 (CBD, 2022). As the recognition of the role IPLCs play in safeguarding biodiversity in Mexico continues to grow, further empirical research that evaluates the conservation potential of these diverse land management types in Mexico and beyond is warranted. Spatial evaluations of the conservation potential for threatened species in territories under Indigenous stewardship are scarce in the Neotropics, and for OECMs and other types of community-led conservation efforts, they are entirely lacking.

**Table 1.**
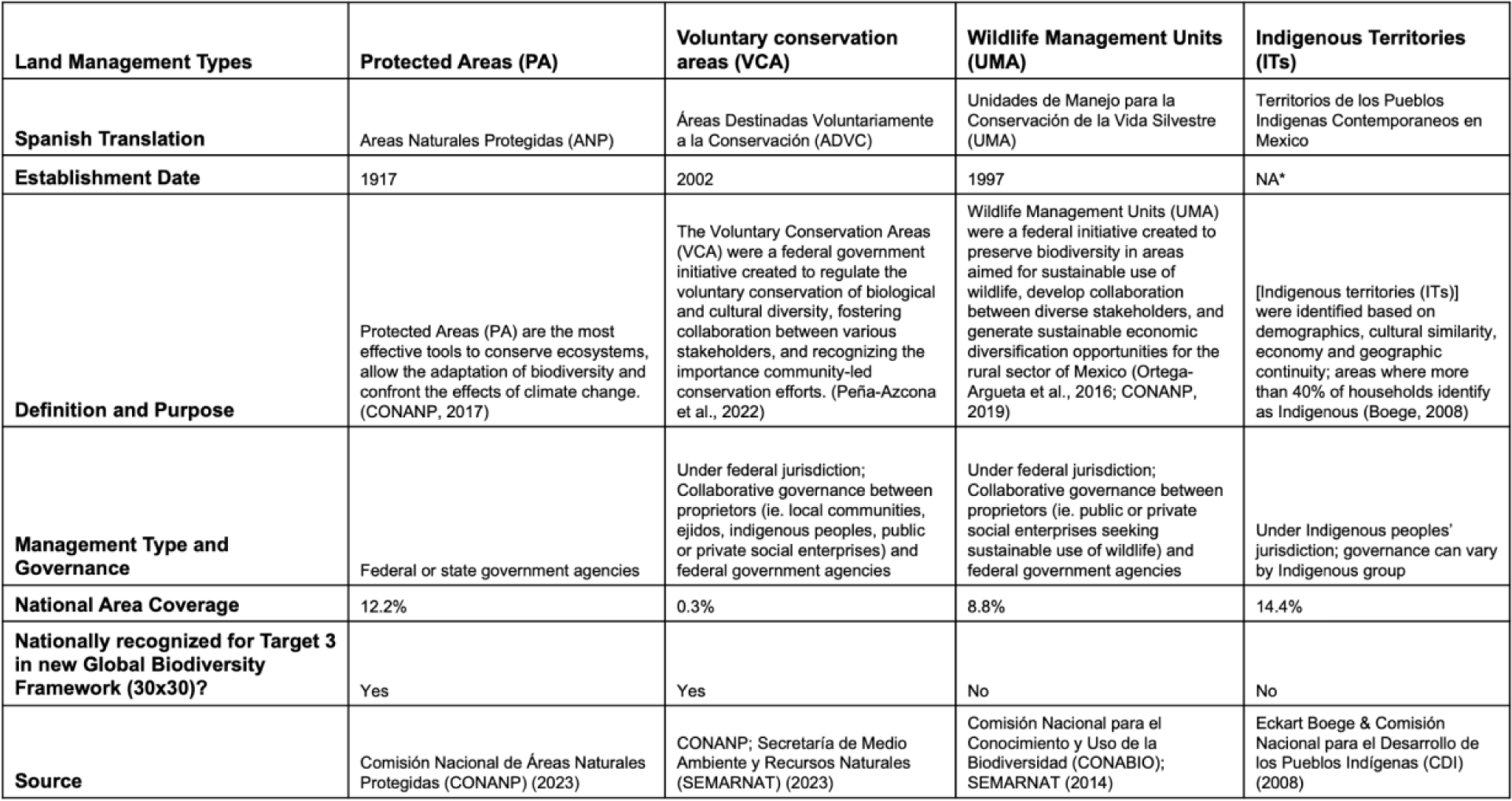
- Different land management types being analyzed in the geospatial analysis. *Note - Recognition of Indigenous territories and rights in Mexico has been an ongoing process marked by historical injustices, legal reforms, and social struggles. It’s important to note that challenges and issues related to Indigenous land rights continue to be a significant topic in Mexican politics and society. Thus, the specific year in which Indigenous lands were officially recognized can vary depending on the region and the historical context.

Here, we conduct a geospatial analysis to evaluate the conservation potential of four Mexican land management types by 1) calculating overlap occurrence between land management types and carnivore ranges, 2) quantifying the distribution of land management types across 29 individual carnivore ranges, and 3) identifying protection gap hotspots where protection expansions could be warranted. We hypothesized that carnivore richness and overlap percentage with carnivore ranges would be highest in PAs and lowest in UMAs based on national designations and coverage area. We expected PAs to be the most strategically placed for carnivore conservation. We also expected extreme heterogeneity in the distribution and co-occurrence of these land management types across the range of carnivores. Finally, we expected protection gap hotspots to be located near existing area-based approaches assuming such areas have been strategically placed to maximize biodiversity protection.

## Methods

### Study area

Our study area comprised the entirety of Mexico (1,972,550 km²), the fourth most biodiverse country in Latin America. In the northern regions, arid climates support desert and grassland ecosystems, while in the southern regions, tropical climates support a mix of rainforest, mangrove, and tropical deciduous forest ecosystems. The presence of the Sierra Madre Mountain range generates unique, specialized ecosystems such as coniferous forests in the northern regions and tropical montane cloud forests in the southern regions of the country. Mexico hosts a large diversity of endangered and endemic carnivores, ranging from the jaguar (*Panthera onca)* to the pygmy raccoon (*Procyon pygmaeus*). Aside from its high biodiversity profile, Mexico is ranked second in endemic languages globally (Otegui-Acha, 2010), with over 65 different Indigenous groups, mostly found in the southern regions of the country (CDI, 2015). This combination of both biological and cultural diversity makes Mexico an ideal system to investigate the complexity of the social-ecological interactions that heavily influence a landscape of varied conservation approaches, governance structures, and stakeholder involvement.

### Geographic information and species range data

We derived national boundaries for Mexico from the Global Administrative Areas database (http://gadm.org/, v.4.1, accessed 2023-03-10) along with four terrestrial conservation area types from various sources (Table 1). These spatial datasets were used to explore the diversity of area-based conservation strategies and the potential for carnivore conservation throughout Mexico. We included ITs that had been recognized by the National Commission for the Development of Indigenous Peoples (CDI) in a collaborative delimitation of Indigenous Territories (Boege, 2008), covering 14.4% of Mexico. We acknowledge that not all ITs have been formally recognized and absences on our map do not indicate areas devoid of the culture, influence, and values of Indigenous peoples. We also included three distinct area-based conservation approaches in Mexico (PA, VCA, UMA). Excluding wetlands and marine-based protected areas (eg. Ramsar sites), terrestrial PAs cover 12.2% of Mexico. UMAs are not formally designated as conservation areas by the Mexican government, covering 8.8% of Mexico. VCAs also contribute to the designated protected estate, covering 0.8% of Mexico. Given the diversity of spaces with heightened conservation potential and their differences in governance and purpose, we hereafter collectively refer to UMAs, PAs, ITs, and VCAs as “land management types” throughout our manuscript.

We obtained a species list from the International Union for Conservation of Nature (IUCN) Red List of Mesoamerican carnivores (IUCN, 2023) and extracted their corresponding extant species ranges maps. Species whose ranges did not occur in Mexico, were not completely terrestrial, or were not in the Order Carnivora were removed. The harbor seal (*Phoca vitulina)*, North American otter (*Lontra canadensis*), and neotropical otter (*Lontra longicaudis*) were excluded because they are not considered terrestrial carnivores. The Mexican gray wolf (*Canis lupus baileyi*) and the brown bear (*Ursus arctos*) were also excluded because these species are considered extinct in Mexico (Miller, 2006; National Academies of Sciences Engineering and Medicine, 2019). This yielded 29 mammalian carnivore species that were included in our analyses (Table S1). Despite some limitations and widespread critiques (Hughes et al., 2021), the IUCN Red List range maps remain a widely-used tool in species distribution models and spatial analyses. However, to confirm the presence of all 29 species within our study extent, we verified occurrences for each species using the Global Biodiversity Information Facility (GBIF) database (www.gbif.org, v.4.1, accessed 2023-05-21).

### Spatial relationships between land management types and carnivore ranges

For all 29 carnivore species, we quantified areas of overlap between ranges and the four selected land management types in Mexico by calculating the area of the spatial intersection polygon. We used this areal calculation to further calculate two overlap percentages in ArcGIS Pro (v.10.8.1, ESRI Inc.)— one proportional to each carnivore range and one proportional to each land management type. Overlap percentages proportional to carnivore range area were informative for understanding how much of a carnivore range was protected by a specific land management type. Conversely, overlap percentages proportional to the area of a land management type revealed how much of a land management type was dedicated to protecting specific species and whether a land management type was considered strategically placed. The latter overlap percentage helped normalize protection coverage, and overall conservation potential, per land management type by total area coverage given the high variability in size across selected land management types. For example, VCAs had a significantly smaller extent compared to ITs or PAs. Thus, having a metric to normalize by size was essential for making informative and holistic conservation potential comparisons between all four land management types. Finally, we also calculated the average for both overlap percentages, proportional to both carnivore range area and land management type area, across all 29 selected carnivores.

We identified which land management types had the highest levels of carnivore richness and highest range overlap percentages. Aside from differences in overall species richness and overlap percentages, we also assessed which land management types had greater overlap percentages with species considered threatened based on IUCN Red List conservation designations (IUCN, 2023), national conservation designations (SEMARNAT, 2010), and with species considered endemic to Mexico. The only endemic carnivores included in our study were the pygmy raccoon and the pygmy spotted skunk (*Spilogale pygmaea)*. We further investigated the relationship between national conservation designation and overlap percentages to test whether a relationship existed between designation level and the protected percentage of carnivore ranges by running a generalized linear model (GLM) analysis using the R statistical software (R Core Team, 2023). For this model, threatened status was selected as the explanatory variable, and total unprotected range percentage was the response variable. No transformations or random effects were incorporated into the model.

To determine which carnivores that had the highest level of land management type co-occurrence within their range, we calculated an overlap occurrence metric. To calculate this metric, we first calculated the overlap percentage total across all four land management types (including non-designated UMA and IT estates) for each carnivore, without accounting for existing overlap amongst land management types. Land management types were then merged into a single polygon and the overlap percentage total for each carnivore with the entire Mexican protection estate was calculated once again to account for the overlap occurring between land management types. The difference between these two total overlap percentages yielded our overlap occurrence metric.

### Gap assessment

To visualize the distribution of diverse land management types inside carnivore ranges, we created individual heat maps that quantified land management type overlap within all 29 carnivore species’ ranges. Carnivore ranges were rasterized and assigned a value of 0. We also rasterized all four land management type polygons and assigned a value of 1. For each carnivore species, we summed land management type and carnivore range raster with their assigned values, resulting in individual species heat maps where co-occurring land management types ranged from 0 to 4 where a 4 indicated an area where all four land management types overlapped within the species’ range.

In addition to creating an individual heat map for each carnivore range, we also extracted areas that had no overlap with any of the selected land management types, defined as protection-absent areas, and aggregated them to create a country-level heatmap (Figure 1). This map revealed protection gap hotspots where additional area-based conservation approaches are needed to increase carnivore conservation potential across the country. To systematically identify regions deemed as protection gap hotspots, we selected areas that had several overlapping protection-absent areas above the 90th percentile.

**Figure 1.**
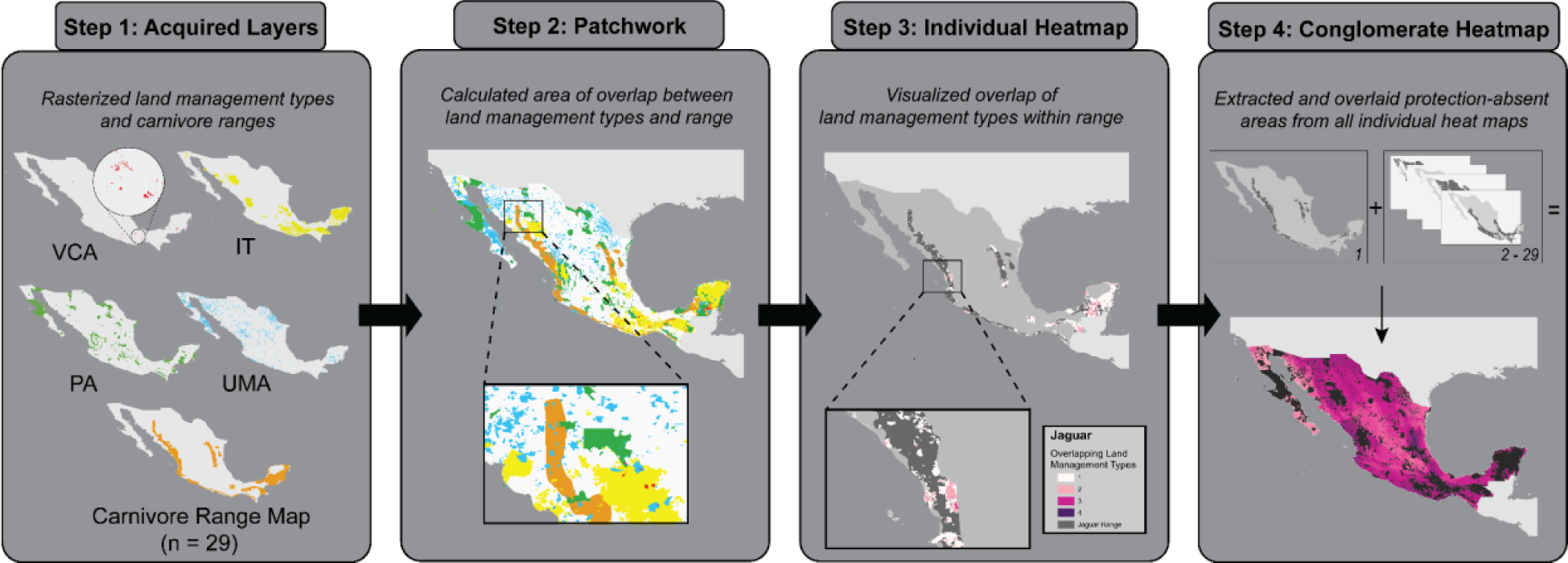
- Workflow of geospatial analysis conducted. We use the jaguar (*Panthera onca)* range (in orange) as an example throughout the workflow.

## Results

### Carnivore conservation potential

We observed minimal variation in species richness across land management types. Overlap percentages, however, revealed major differences between the four land management types and their conservation potential (Appendix S1). The PA estate demonstrated substantial variability in its percentage of overlap (proportional to carnivore range) with carnivore ranges, ranging from 5.5% (*Spilogale pygmaea*) to 55.2% (*Procyon pygmaeus*). On average, 12.8% of the PA estate overlapped with carnivore ranges. On average, the IT estate exhibited an overlap percentage of 19.16%, with carnivore ranges ranging from 0.008% (*Spilogale putorius)* to 45.95% (*Bassariscus sumichrasti*) in percentage of overlap. The UMA estate exhibited a higher average overlap percentage with carnivore ranges (7.29%) than the VCA estate (0.44%). The UMA estate varied from 2.45% (*Spilogale angustifrons*) to 14.13% (*Vulpes macrotis*) in overlap with carnivore ranges, while the VCA estate varied from 0.003% (*Spilogale putorius*) to 0.98% (*Bassariscus sumichrasti*). Calculations of spatial overlap were also determined proportionally to the total land management type area, which ranged 42.2% (UMAs) to 53.9% (VCAs). For overlap percentage proportional to land management type which assessed strategic placement, carnivore ranges overlapped with 53.9% of the VCA estate (Range: 0 to 99.79%), 52.3% of the IT estate (Range: 0 to 99.17%), 43.5% of the PA estate (Range: 0.11 to 97.64%), and 42.2% of the UMA estate (Range: 0 to 98.92%) on average, making VCAs the most strategically placed for carnivore conservation out of selected land management types. For individual carnivore ranges, we also found three cases where specific land management types were essential in providing coverage for carnivore species (Figure 2).

**Figure 2.**
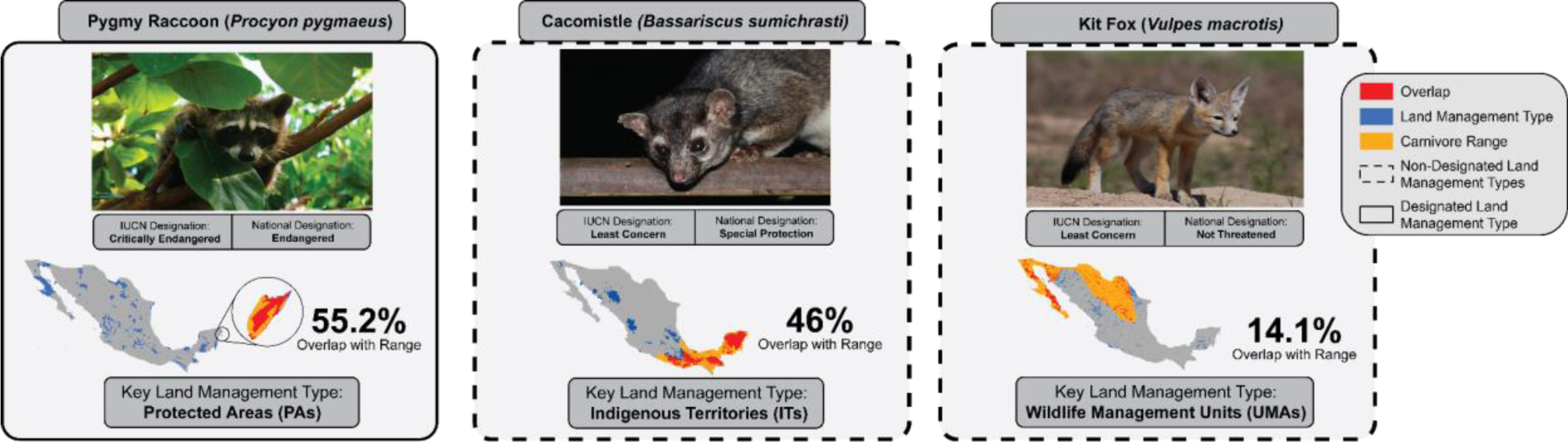
- Case-studies for three species and their “key” land management types. Key land management types were selected per species based on overlap percentage. *photo credits: Laura Gaudette (Bassariscus sumichrasti), Christopher Gonzalez (Procyon pygmaeus), & James Marvin Phelp (Vulpes macrotis)*.

Under the officially designated protection estate in Mexico (PAs and VCAs), only one species (*Procyon pygmaeus)* had more than 50% of its range protected. On average, 87% of carnivore ranges were without any protection (Figure 3a). Including non-designated land management types (UMAs and ITs) yielded only six carnivores that had more than 50% of their range protected: the jaguar, the southern spotted skunk (*Spilogale angustifrons*), kinkajou (*Potos flavus*), cacomistle (*Bassariscus sumichrasti)*, striped hog-nosed skunk (*Conepatus semistriatus*), and pygmy raccoon (Figure 3b). On average, 65.2% of carnivore ranges were without any protection including non-designated land management types.

**Figure 3.**
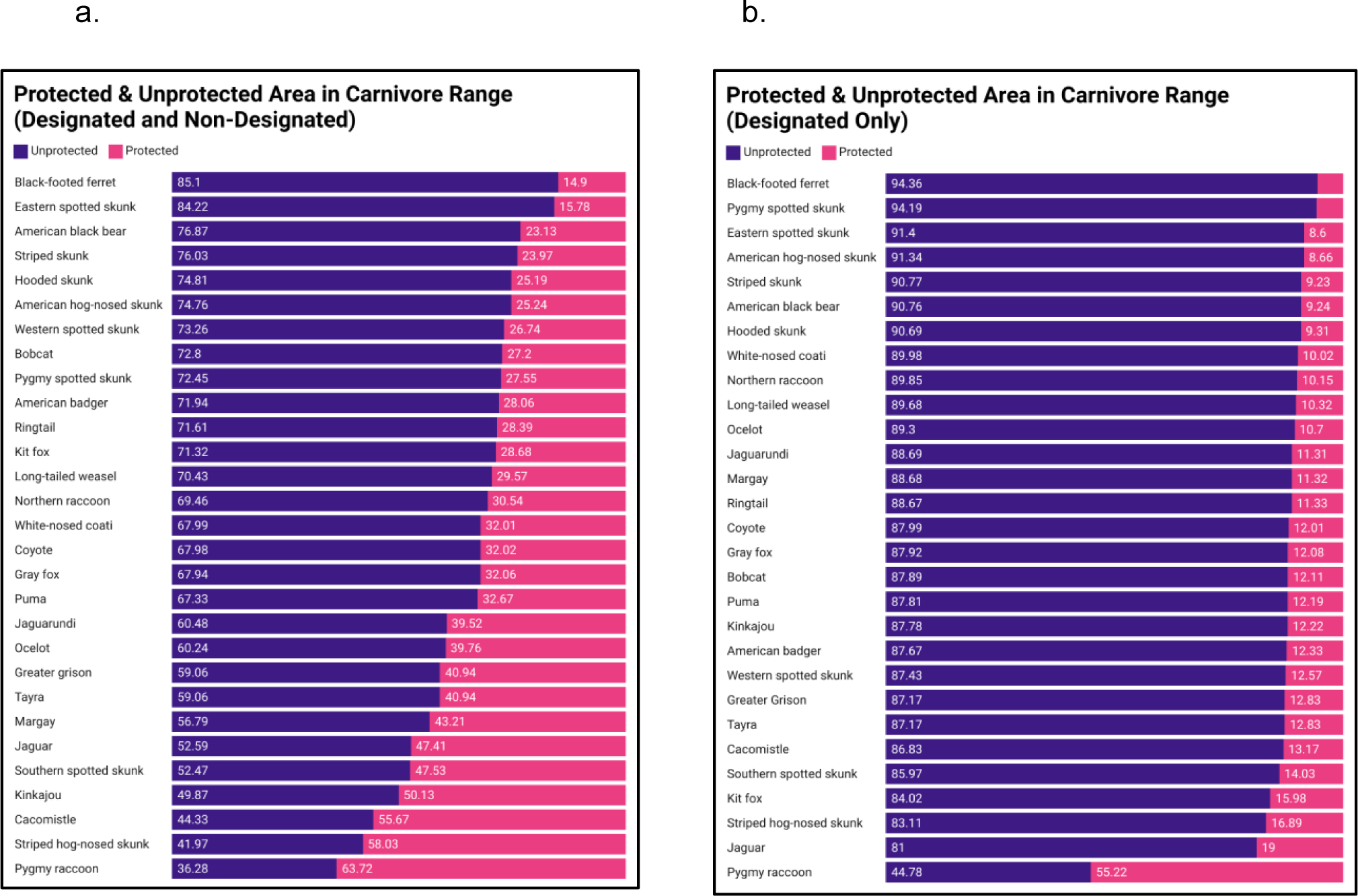
- Coverage of protected areas within the ranges of 29 Mexican carnivores: a) protected area includes only lands officially designated by the Mexican government that includes protected areas and VCAs, b) all land management types included in protected area category. *Bar graphs were created using Datawrapper tool (*https://www.datawrapper.de/)

### Threatened and endemic carnivores

Our results revealed differences between national and IUCN threatened designations in respect to land management type coverage (including both designated and non-designated). We found that threatened carnivores with an IUCN designation of either “Vulnerable”, “Endangered”, or “Critically Endangered” (n = 4) averaged 69.5% unprotected area within their ranges. The black-footed ferret (*Mustela nigripes*; Endangered) was perhaps the most vulnerable species of this group, with 84% of its range unprotected, followed by the eastern spotted skunk (*Spilogale putorius;* Vulnerable), with 84.2%, the pygmy spotted skunk (Vulnerable), with 72.5%, and the pygmy raccoon (Critically Endangered), with 36.3%. Conversely, we found that carnivores with a Mexican conservation designation of “Special Protection”, “Threatened” and “Endangered” (n = 15) averaged 58.8% of unprotected area within their ranges, compared to an average of 72% unprotected area for non-threatened carnivores (n = 14). Species considered “Special Protection” averaged 45.4% of unprotected area (n = 3), followed by species considered “Threatened” (n = 6; 67.2% unprotected), and species considered “Endangered” (n = 5; 57% unprotected). Carnivores endemic to Mexico (n = 2), averaged 54.4% of unprotected area within their ranges, with the pygmy raccoon having 36.3% of its range unprotected and the pygmy spotted skunk having 72.5% unprotected.

Specific land management types offered significant coverage for some threatened carnivores. For overlap percentages proportional to range, we found that the PA estate provided the highest level of protection coverage for the four IUCN threatened carnivores, with an average overlap of 18.8% with their ranges. The PA estate also provided the highest level of protection coverage for threatened carnivores in Mexico under national conservation designations (n = 15), with these carnivore ranges overlapping with 25.54% of the PA estate on average, followed by the IT estate at 14.8%. For endemic carnivores, the PA estate contributed the highest level of protection coverage for the pygmy raccoon with an overlap percentage of 55.2%, while the IT estate contributed the highest level of protection coverage for the pygmy spotted skunk (17%). For overlap percentages proportional to land management type area, we found that VCAs and ITs were the most strategically placed for threatened carnivore conservation. Ranges for carnivores considered threatened under Mexican designation had the highest overlap occurrence with both the IT and VCA estates, with an average overlap percentage of 52.7% and 49.7%, respectively. Our GLM analysis revealed that having a threatened designation of “Special Protection” or “Threatened” under Mexican designation did not have a significant effect on the amount of total protection found inside a carnivore range (p = 0.07, p = 0.05, respectively). However, having a “Not Threatened” or “Endangered” designation significantly influenced the amount of total protection found inside a carnivore range (p < 0.002, p < 0.0001, respectively). For IUCN designations, we found that “Critically Endangered” (p < 0.0001), “Endangered” (p <0.01), “Vulnerable” (p < 0.002), and “Least Concern” (p < 0.01) designations all significantly influenced the amount of total protection. Only the “Near Threatened” (p = 0.14) designation had no significant effect on the amount of total protection found inside a carnivore range.

The overlap occurrence metric measuring co-occurrence of land management types within carnivore ranges ranged from 0.41 (eastern spotted skunk) to 9.11 (pygmy spotted skunk). Four species were selected as having an overlap occurrence metric in the 90th percentile, meaning that overlap amongst land management types was highest within their ranges. These species included the jaguar (8.53), cacomistle (*Bassariscus sumichrasti,* 6.97), pygmy raccoon (9.11), and the striped hog-nosed skunk (8.10), all of which are considered threatened under Mexican conservation designations.

### Gap assessment

Individual heat maps revealed areas with high conservation potential due to the absence of any protective land management type (Appendix S2-30). In most cases, Mexican carnivores did not have areas where all four land management types overlapped within their range. By aggregating areas with no protective land management for individual species, the resultant conglomerate heat map highlighted potential areas for future protection (Figure 4). We identified five areas as protection gap hotspots where area-based conservation approaches could be warranted in the future. One protection gap hotspot was in the state of Chihuahua in a desert ecoregion at the United States-Mexico border, where 17 carnivore protection-absent areas overlapped, though majority of the species found in this hotspot were designated as “Not Threatened” by the Mexican government. The two hotspots on the western coasts, in the states of Nayarit (n = 20) and Guerrero (n = 19), had higher carnivore richness than the Chihuahuan hotspot and slightly higher levels of threatened carnivores. We found the hotspots on the eastern coast, in the states of San Luis Potosi and Veracruz (n = 22) and southern Veracruz (n = 20), to be the most critical for conservation expansion since they had a higher carnivore richness, and more than half the carnivore species were of conservation concern.

**Figure 4.**
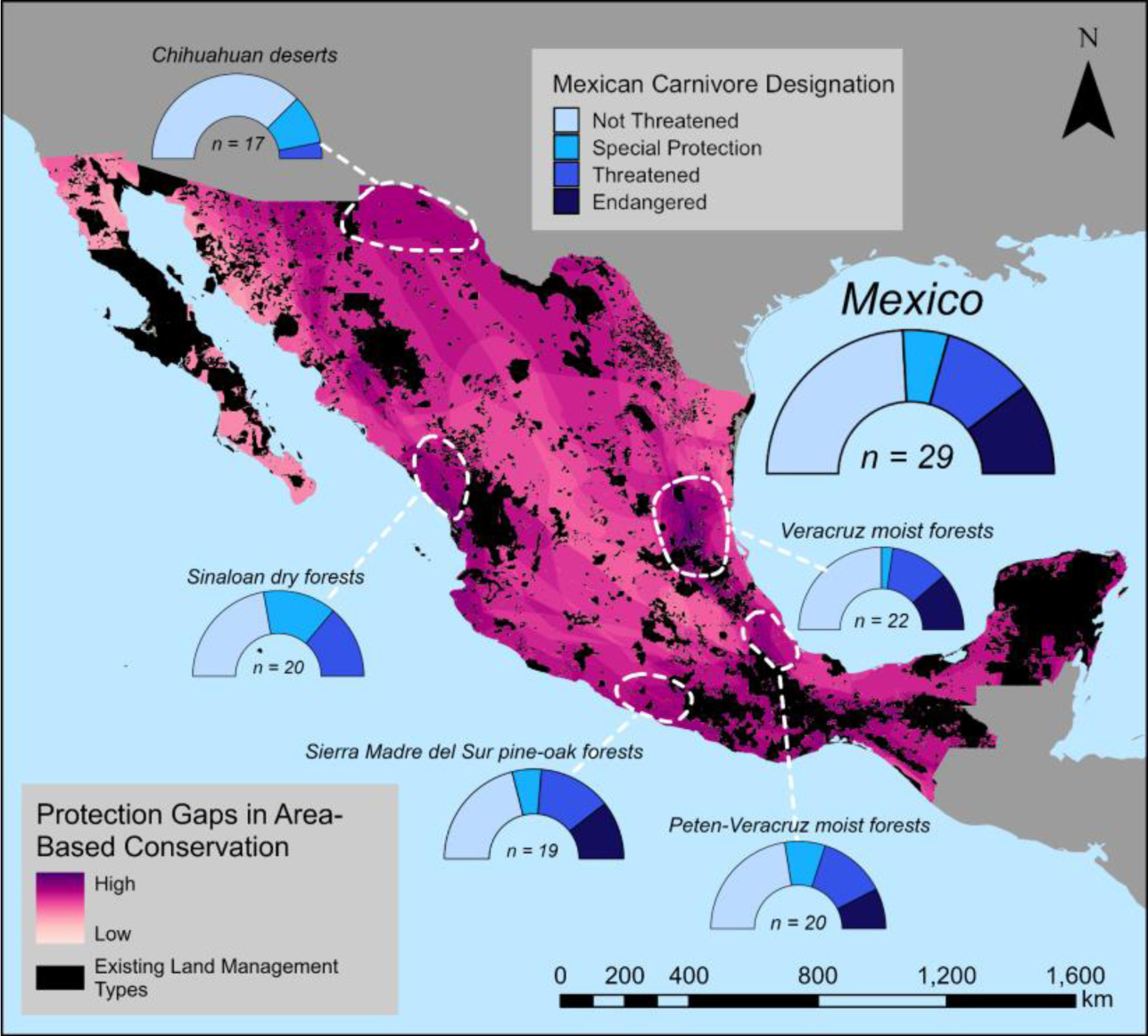
- Conglomerate heat map made up of overlaid protection-absent areas from 29 species, highlighting areas where there are gaps in area-based protection estate. Areas in black are the selected land management types (ITs, PAs, VCAs, and UMAs). Areas in light pink are regions where fewer protection-absent areas overlap, indicating lower species richness, and low priority gaps for area-based conservation measures. Areas in purple are regions where many protection-absent areas overlap, indicating high priority gaps in area-based conservation measures. Areas circled in white are protection gap hotspots, and callouts indicate the predominant ecoregion and Mexican conservation designation breakdown of all carnivore ranges occurring inside hotspots. *Created using ArcGIS Pro (v.10.8.1, ESRI Inc.)*.

## Discussion

With the heightened international momentum to reach post-2020 biodiversity targets by 2030, countries are now presented with the logistical challenges of meeting these conservation goals and ensuring that they are effectively implemented (Butchart et al., 2014; Saura et al., 2018). One such target centers on expanding the conservation estate. Our study complements ongoing expansion conversations by exploring the current role of Mexico’s heterogeneous land management systems in protecting terrestrial carnivores and recommending new protected areas. We examined spatial co-occurrence patterns of 29 terrestrial carnivores in relation to overlap with various forms of land protections across Mexico. We found that 65.15% of carnivore ranges on average remain unprotected in Mexico, even with the inclusion of non-designated land management types (ITs and UMAs). With mounting concerns of carnivore biodiversity loss, we also closely evaluated threatened carnivores (by IUCN designation, national designation, and endemic status) in our spatial analysis, and found that dour threatened species had the highest levels of land management type co-occurrence (above the 90th percentile) in their ranges, indicating the need for strategic conservation planning to facilitate communication amongst diverse stakeholders. Five protection gap hotspots— areas where there is no existing protection through explicitly delimited designated or non-designated conservation spaces— were identified across a variety of Mexican ecoregions, suggesting where conservation estate expansions may be warranted to reach biodiversity targets.

Mexico’s terrestrial protected area coverage currently sits at 14.6%, with an additional 15.4% needed to reach 30% protection coverage as set by international biodiversity targets. Our geospatial analyses provided an optimistic lens into the conservation of carnivore populations in Mexico based on the coverage and co-occurrence of four selected land management types, and their collective potential to contribute to biodiversity targets. VCAs may be most strategically placed, with the highest average range overlap (proportional to VCA area) across all carnivore species ranges (53%), and with the second highest average overlap with nationally designated threatened carnivore ranges. This was followed closely by the IT estate, suggesting that areas managed by IPLCs have critical spatial potential for carnivore conservation. Though VCAs, and by extension UMAs, are smaller units and may alone lack the capacity to solely offer protection for larger-bodied carnivores that exhibit large home ranges (Ripple et al., 2014), they can help reinforce connectivity when located near adjacent PAs or when linked with wildlife corridors (Saura et al., 2019). Moreover, these area-based conservation approaches are exemplary in centering proprietors’ needs through a shared governance structure to reach biodiversity targets, which support the departure from conventional conservation strategies and paradigm shift towards a more inclusive and effective conservation methodologies (Vazquez-Quesada & Jiménez, 2020; Peña-Azcona et al., 2022). UMAs, for example, allow for traditional hunting efforts by local communities to continue while simultaneously implementing local biodiversity targets (Ortega-Argueta et al., 2016). Our spatial analysis is supported by many studies that highlight important overlaps between ITs, community-conserved areas, or OECMs, and federal PAs for a variety of focal taxa. For example, evidence for the spatial importance of ITs, community-conserved areas, or OECMs through spatial co-occurrence analyses has been found for bats in the Amazonian basin (Fernandez-Llamazares et al., 2021), vertebrates in Brazil, Canada and Australia (Schuster et al., 2019), primates in the Neotropics (Estrada et al., 2022), jaguars in Latin America (Figel et al., 2011; Figel et al., 2022), and tropical forests in Latin America (Bray et al., 2008; Qin et al., 2022). Indigenous, or local community-led conservation approaches often contribute to shifts in local community attitudes towards conservation efforts (Störmer et al., 2019; Everard et al., 2016; Vimal et al., 2021). Moreover, several case studies in Latin America have found that the adoption of successful conservation strategies that yield increases in revenue through ecotourism, government payment for ecosystem services (PES) programs, and sustainable resource extraction in a community, can produce a domino effect on surrounding communities by encouraging them to also engage in conservation efforts (Berkes 2008; Bray et al., 2012; Premauer & Berkes, 2015; Blackman 2015). From an expansion perspective, the benefits of encouraging similar OECMs with collaborative governance structures are quite clear, and supporting these kinds of conservation efforts may be the solution to meeting expansion targets by 2030.

Our overlap occurrence metrics calculated for all carnivores underscore the opportunities for transfers of knowledge, co-management strategies, and inclusive conservation strategies that result from supporting diverse mosaic of land management types for carnivore conservation (Fitzsimons & Wescott, 2008; Velho et al., 2016; Archibald et al, 2020; Vimal et al., 2021). It is well understood that conservation management is most successful when there are attempts at collective actions that involve all stakeholders in spaces where overlapping governance structures are already present (Bray et al., 2012). However, working with co-governing or in multi-governance spaces can also be complex given the intricacies of a multistakeholder reality, ranging in environmental value systems amongst different communities (Jones et al., 2016) and the existing management tensions that result from stakeholders’ varying needs (Raymond et al., 2022). Others have discussed the wide range of economic outcomes that can be produced by different area-based conservation measures with varying governance structures related to wildlife ecotourism, which can also be a point of contention (Chidakel & Child, 2022). For example, the development of multiple governance structures occurring within a protected area in transfrontier conservation areas like the Great Limpopo Transfrontier Park in Southern Africa have created specific place-based challenges that require conservation effectiveness analyses at more local scales (Mpofu et al., 2023). In the context of Mexico, multi-stakeholder collaboration in these areas is further hindered through the existing power dynamics that place pressure on IPLCs to follow government-designed schemas, and the multi-governance schemes already existing even within OECMs (Peña-Azcona et al., 2018). Evaluations of conservation effectiveness for VCAs and OECMs are already lacking at the national level in Mexico and suggesting the expansion or inclusion of these OECMs into the national estate requires a baseline understanding of their performance (Ortega-Argueta et al., 2016; Peña-Azcona et al., 2022). Spatial coverage and overlap of the protection estate alone, while important for determining conservation potential, are not enough to make comments on the conservation effectiveness of a conservation area (Brooks et al., 2004). Thus, promoting the proliferation of collaborative governance structures and bottom-up strategies for area-based conservation will require conservation practitioners and managers to think critically about how to measure their conservation effectiveness, including both social and ecological effectiveness indicators, before implementing in-situ conservation in Mexico. We suggest future research towards a standardized understanding of the effectiveness of ITs and OECMs (VCAs and UMAs) in conserving biodiversity, as this will provide insight into implementation of conservation plans, and a pathway towards OECM integration in multiple-use, multi-governance spaces where biodiversity protection is warranted.

Our gap analysis revealed locations in Mexico where protected area expansions may be targeted. Gap analyses remain a key geospatial planning tool to guide conservation efforts (e.g., Harris et al. 2023). In evaluating the coverage of landscape management types with the distribution of carnivore richness, we identified five protection gaps for carnivores in a range of ecoregions in Mexico. The two protection gap hotspots located on the eastern coasts of Mexico contained the highest level of carnivore richness and a higher number of carnivores considered “Threatened” or “Endangered” under Mexican designation. Thus, we consider these hotspots to be the highest in priority for carnivore conservation efforts. Our study’s identification of hotspots in the state of Chihuahua at the US-Mexico border and on the northeastern coast in the state of Nayarit complements the prior work of Valenzuela-Galvan et al. (2007) and Valenzuela-Galvan & Vazquez (2008), who employed human density measurements, existing protected areas (PAs), and carnivore ranges to delineate carnivore conservation priority areas. The hotspot located in the state of Chihuahua at the US-Mexico border is in a historically contentious space where militarization and conservation have co-evolved, which presents environmental challenges in a hybrid, multi-purpose landscape seeking to protect nature and national security (Meierotto, 2014). Three of our protection gap hotspots (e.g., Veracruz, Guerrero) are located near operational zones for cartels, and traverse many known narcotic trafficking routes (Henkin, 2020). Clashes between area-based conservation efforts and national armed conflicts are becoming increasingly informative for determining PA placement (Hsiao et al., 2023). The range of determined protection gap hotspot locations emphasize the importance of recognizing the overarching social and political dynamics that exist throughout Mexico, which may hinder the expansion of the conservation estate.

In working with spatial, two-dimensional data in Mexico, limitations may exist due to the scale of the analysis, the unavailability of certain spatial data, and the differences in which explicitly-spatial data are available. The variety of spatially explicit data used for land management types and carnivore ranges present some challenges. For example, ITs produced by Boege & CDI (2008) were different from Indigenous land shapefiles found on other databases (ie., LandMark). This variability in Indigenous land or territory boundaries in Mexico serves as a reminder of the difficulty that exists in mapping ITs, as there may be due to the differences in national perceptions of Indigeneity across the landscape, or greater socio-political agendas that might serve to either support or undermine IPLC sovereignty by having varying sizes of ITs (Chapin et al., 2005). Moreover, in recognizing the important roles played by Indigenous people in the protection of biodiversity globally, we also acknowledge that Indigenous Territories contribute more than a heightened biodiversity conservation potential, and that an existing conservation schema or plan should not be considered compulsory, as this reduces Indigenous agency (Jonas et al., 2017). Indigenous peoples and their lands host a range of socio-cultural, biocultural uniqueness that can serve as a multiple-use landscape model from which future conservation and management strategies could learn. Spatial limitations of using carnivore range maps provided by the IUCN Red List database are also worth acknowledging given that boundaries of population distributions may be rough, inaccurate, and often mismatch with occurrence records provided by other databases (Hughes et al., 2021; Chen et al., 2023; Higino et al., 2023). This is especially true for narrow endemic species with smaller home ranges (e.g., mesocarnivores), who tend to have precise ranges than more wide-ranging species (e.g., large carnivores) (Marsh et al., 2022). However, existing reliable and accurate data for carnivore ranges are unfortunately difficult to obtain and validate, as land-use change and other pressures force dynamic distributions (Di Minin et al., 2016; Leão et al., 2023). Other expert range maps such as the ones harmonized by Marsh et al. (2022) (Burgin et al., 2020; Map of Life (https://mol.org)) are partially complementary to the IUCN range maps, and when comparing our selected carnivores’ ranges between the two set of range maps, we did not find a systematic bias indicating consistent over- or underestimations in comparison to other sources. For example, the IUCN Red List provided a smaller range for the jaguar in comparison to Burgin et al., 2020, which conversely provided a smaller range for the American black bear in comparison to the IUCN Red List. To understand the spatial distribution of carnivore species, combining range maps with accumulating data from camera traps (Chen et al., 2023) or other occurrence records (e.g., GBIF) are warranted for the creation of more precise species range maps and improved conservation mapping and decision mapping.

Moreover, major differences in threatened status between IUCN and Mexican national designations revealed the importance of national designations in providing a more localized, country-level designation that can better inform protected area expansions for locally threatened species. Only two of the 15 carnivore species considered threatened by Mexican conservation designations were also classified as threatened by the IUCN. Our analyses revealed that national threatened designations aligned more closely with the amount and placement of protection granted by diverse land use management types. While certain species may be of least concern at a global scale, subpopulations of species can be threatened at local scales. Similar to other studies, we recommend Red Lists at subglobal scales given the incongruencies in national and IUCN designations, and in protection coverage offered based on designations (Gardenfors, 2001; Miller et al., 2007). Finally, the question of which land management type to expand or select for a protection hotspot gap remains unanswered, and requires a greater understanding of OECM effectiveness, the local socio-ecological contexts, and the potential for co-governance strategies. While this study highlights protection gap hotspots using information from existing protection areas, a more textured, spatial analysis that accounts for the spatial heterogeneity of land-use, forest cover and risk could help refine hotspots and inform prioritization of expansion (Harris et al., 2022). For example, textured range maps with information on available conservation capacity have shown to be more informative in locating where conservation of wildlife may be necessary (Harris et al., 2023). Moreover, spatial evaluations of the cultural and economic importance of carnivores across Mexico may help gauge where VCAs or UMAs could be established to aid in protection expansions and may help create complementary conservation plans.

## Conclusion

Our study is the first to explore the geospatial relationships between protected areas (PAs), Indigenous Territories (ITs), and other effective area-based conservation measures (OECMs) including voluntary conservation areas (VCAs) and wildlife management units (UMAs), and their overlap with carnivore ranges across Mexico. Human-carnivore interactions are increasing along urban-wildland spectra and development gradients, generating a plethora of multiple-use landscape scenarios with heightened heterogeneity at social, political and ecological scales (Su et al., 2022; Schmidt et al., 2023). The decision to include a diverse suite of conservation approaches in our spatial analysis reflects the emerging reality of Mexico, and many other biodiverse spaces around the world. It is also important to note the rarity of “pristine” landscapes in Mexico and other places in the Global South, since even wildland areas are often inhabited, multifunctional landscapes (Butchart et al., 2014; Adams et al., 2023), making it imperative to move forward with conservation planning through a co-governance or multi-governance framework to enhance local participation and produce resilient conservation efforts. With such varied land management types across Mexico, we aimed to identify the synergistic effects of a holistic protection estate rather than elevating one strategy over others. Mexico currently has 14.6% of its land designated as protected. However, across all four selected land management types, there was a total area coverage of 697,113 km² (35.71% of Mexico). With an average of 87% of carnivore territory remaining unprotected under designated-only land management types, reaching post-2020 biodiversity targets set by the Kunming-Montreal Global Biodiversity Framework may require recognizing the contribution of non-traditional conservation approaches like OECMs (UMAs) and ITs for carnivore conservation. Including UMAs and ITs as nationally designated conservation spaces would prevent having to find new land to occupy for new PAs, especially if an existing, conservation-oriented governance framework is already in place. As the conservation field adjusts to this reality, while simultaneously promoting protection estate expansions, understanding the current protection estate and assessing existing conservation potential are essential for determining where expansions should occur, and which expansion pathways are more effective for promoting coexistence efforts. To foster biodiversity protection, particularly for keystone carnivore species, it is imperative to champion expansions while ensuring effective engagement in conservation planning with IPLCs. A justice-oriented approach in implementing these expansions will be crucial for establishing a global protected area network that not only addresses biodiversity loss and mitigates the impacts of climate change but also fosters collaboration among diverse land management efforts, governance structures, and stakeholders.

## Acknowledgements

We would like to express our sincere gratitude to Dr. Alejandro Ortega-Argueta and Lucia Bustos for assisting in the early conceptual developments of the project. We also acknowledge the invaluable support from lab members in the Applied Wildlife Ecology Lab at the Yale School of the Environment with figure development and study design, especially S. Gamez and E. Jones. Finally, we also express gratitude to Indigenous Peoples and rural communities throughout the Mexican landscape that have helped preserve carnivore populations, not only by contributing tangibly to conservation efforts, but also by uplifting the cultural importance and longevity of many carnivore species in Mexico.

## Author Contributions

Gonzalez, Germar: Project Administration, Conceptualization, Data Curation, Methodology, Formal Analysis, Writing - Original Draft; Harris, Nyeema: Supervisor, Conceptualization, Methodology, Writing - Reviewing and Editing, Resources, Funding Acquisition

