## Supplementary figures and images for "Key unprotected areas for carnivore conservation in Mexico"

### AmericanBadgerHM.png

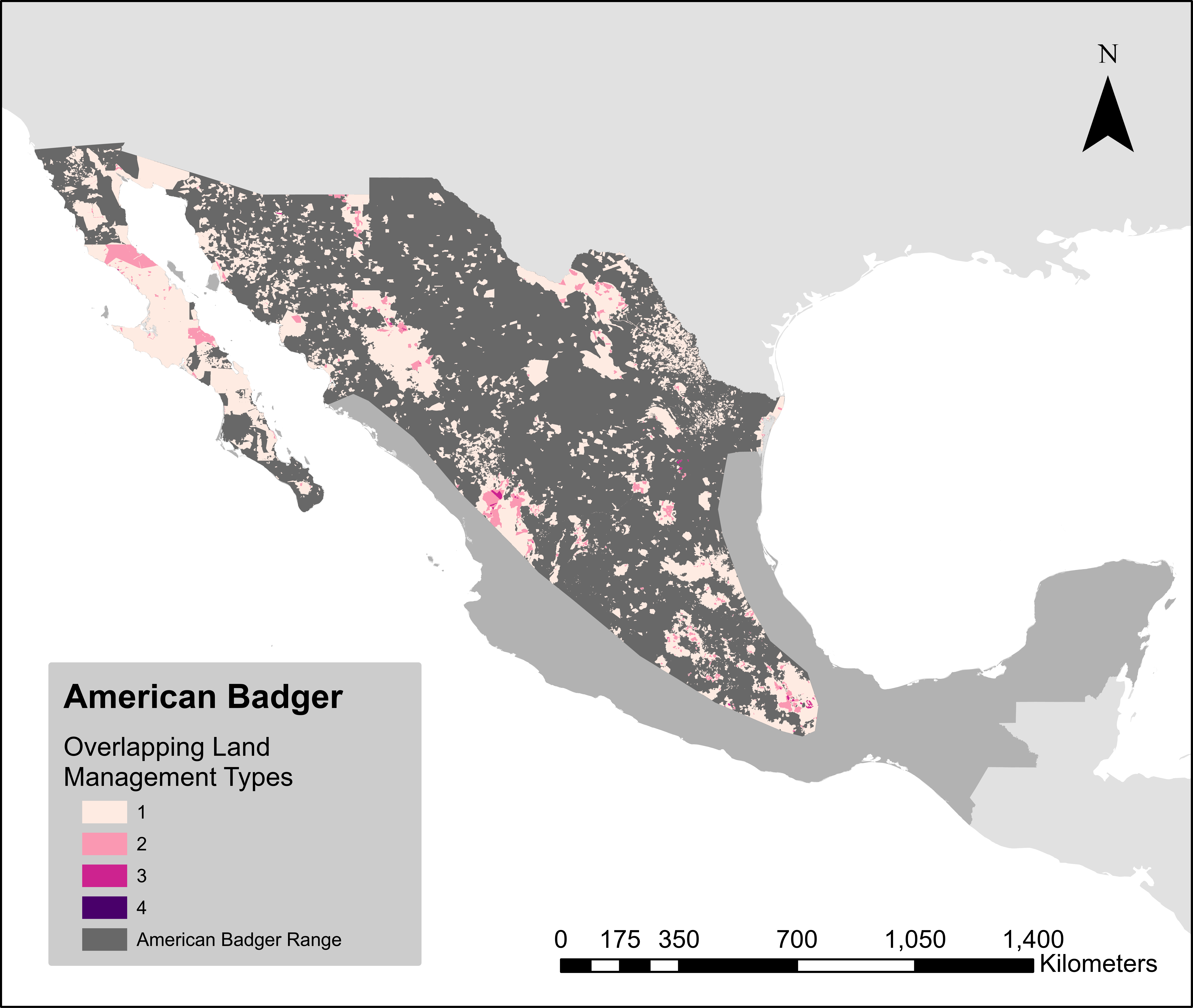

### AmericanBlackBearHM.png

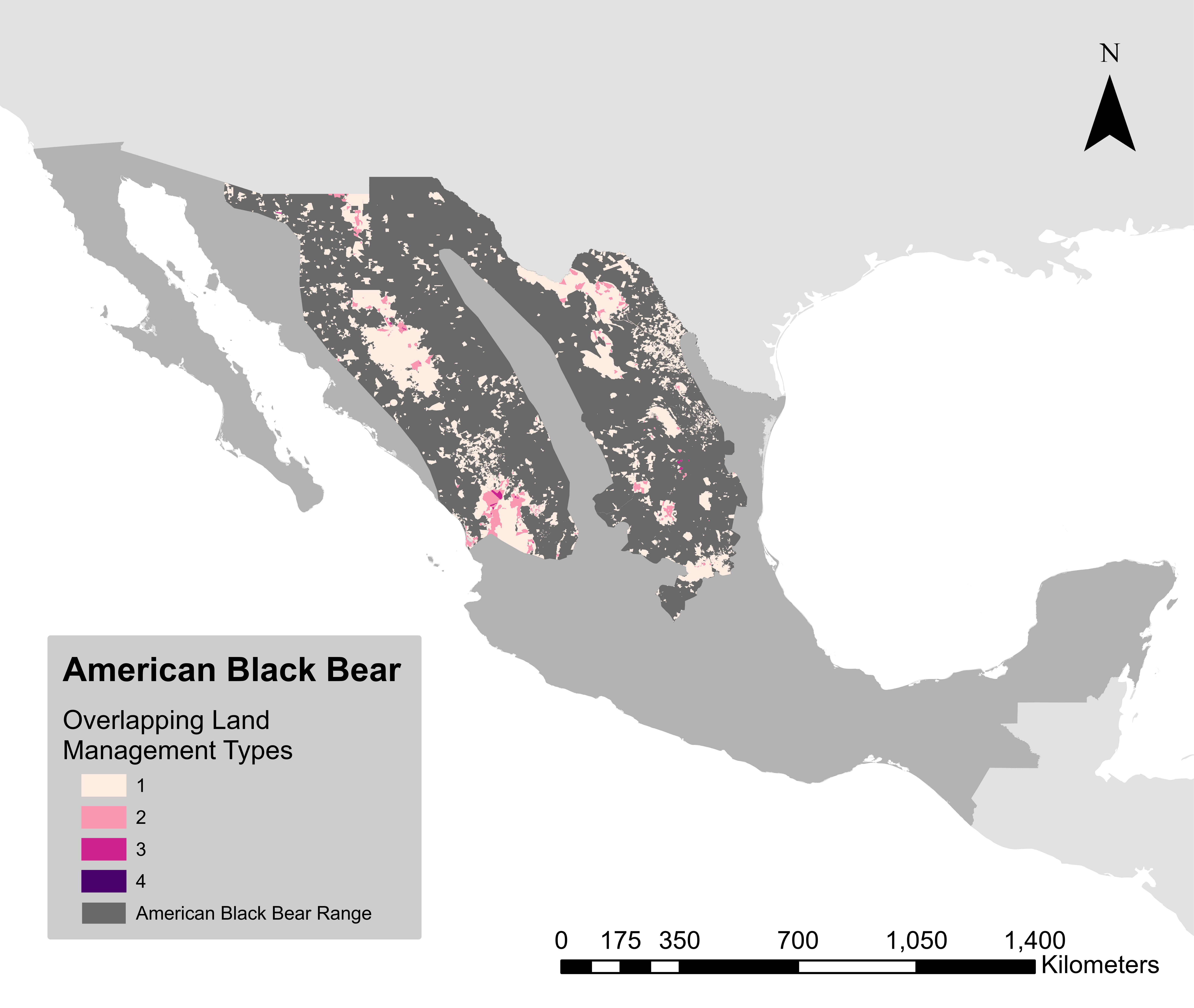

### AmericanHogNosedSkunkHM.png

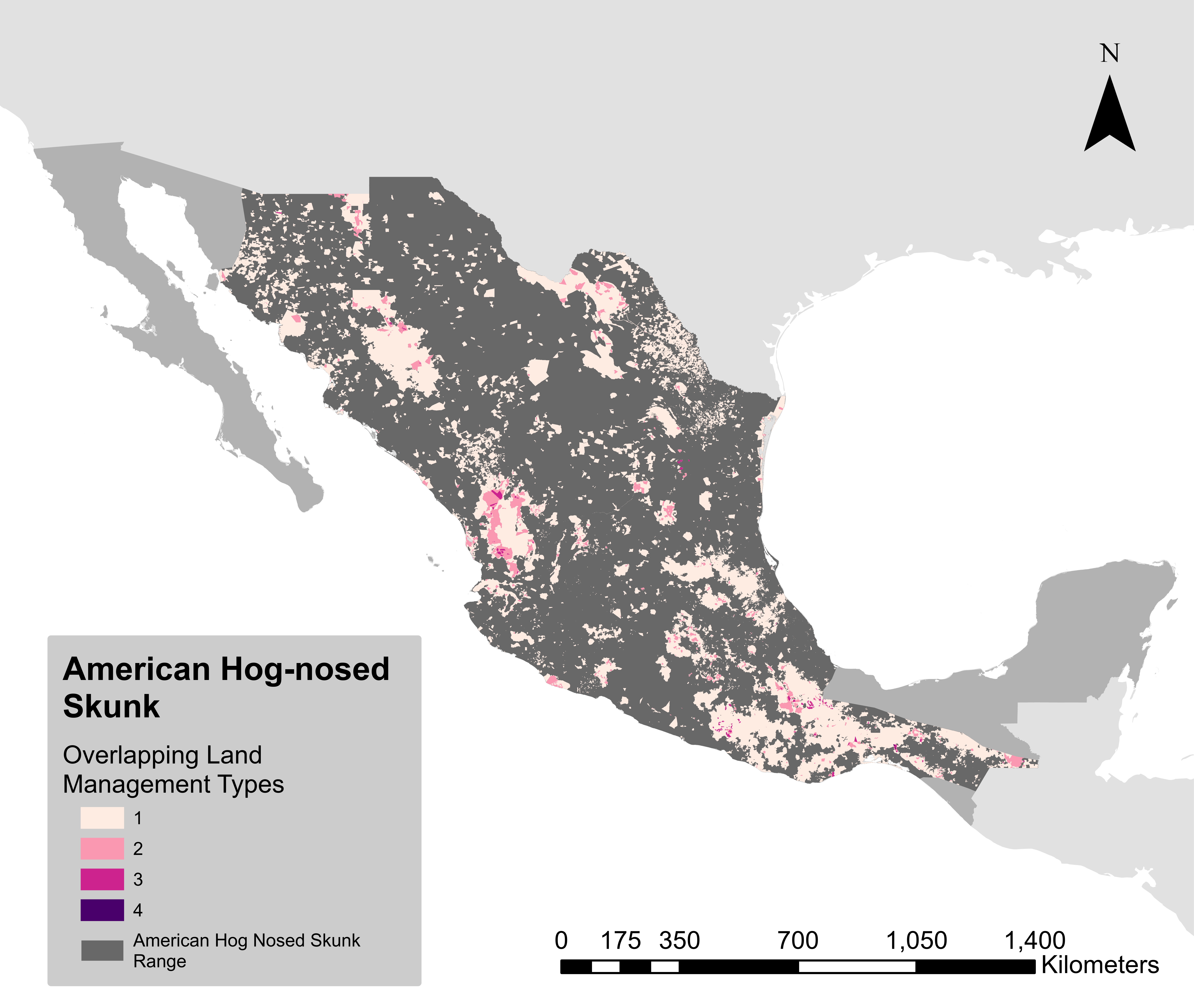

### BlackFootedFerretHM.png

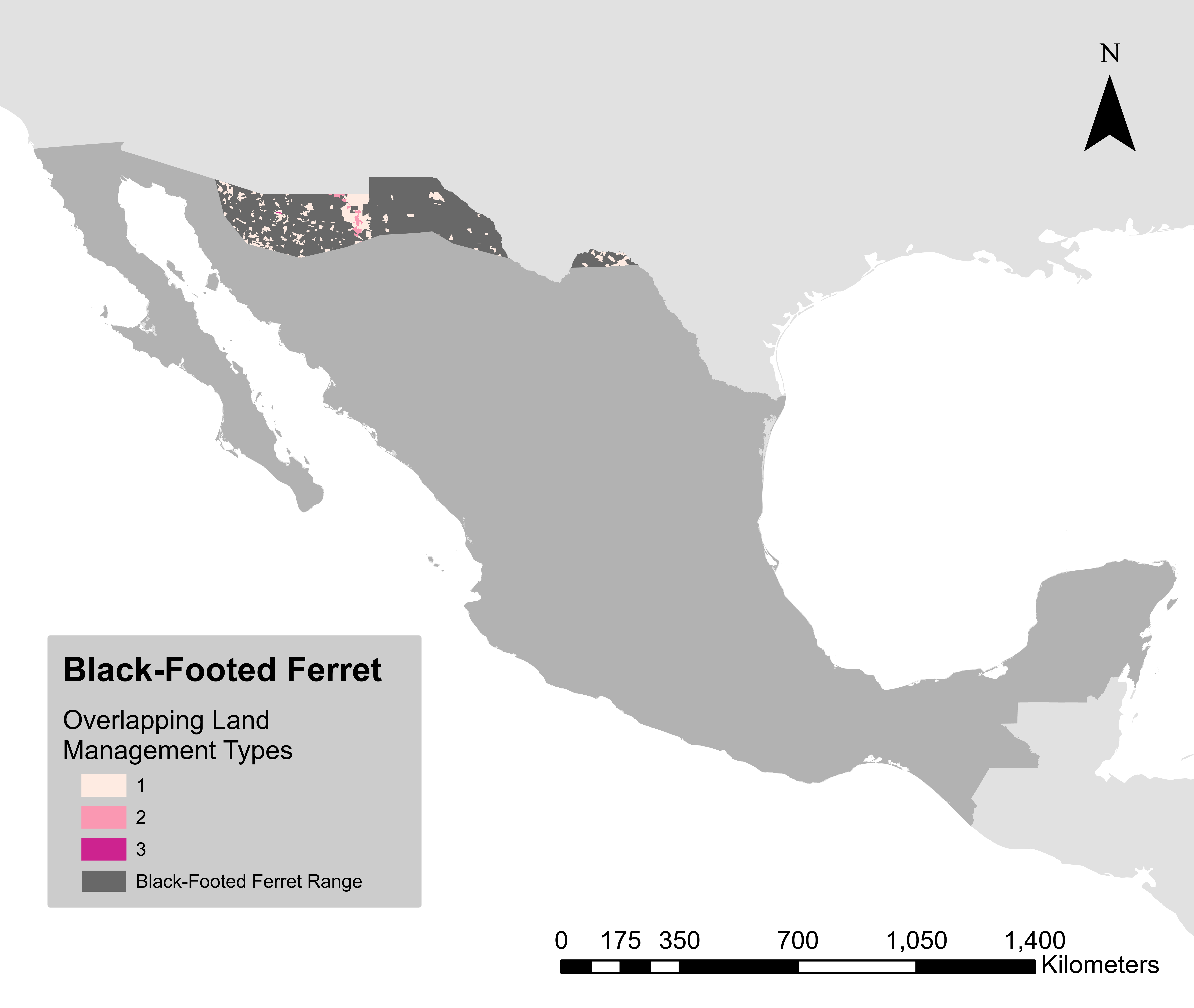

### BobcatHM.png

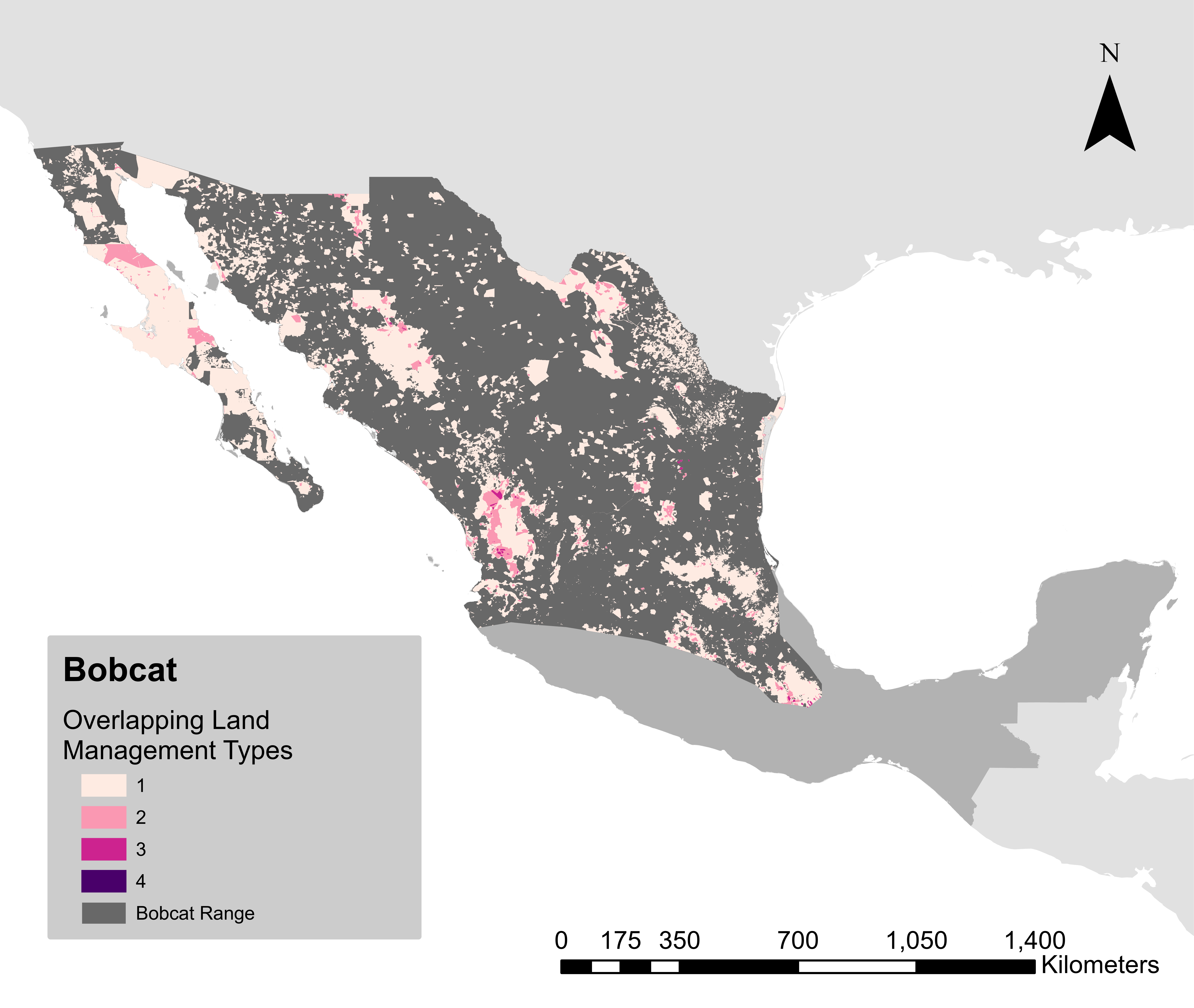

### CacomistleHM.png

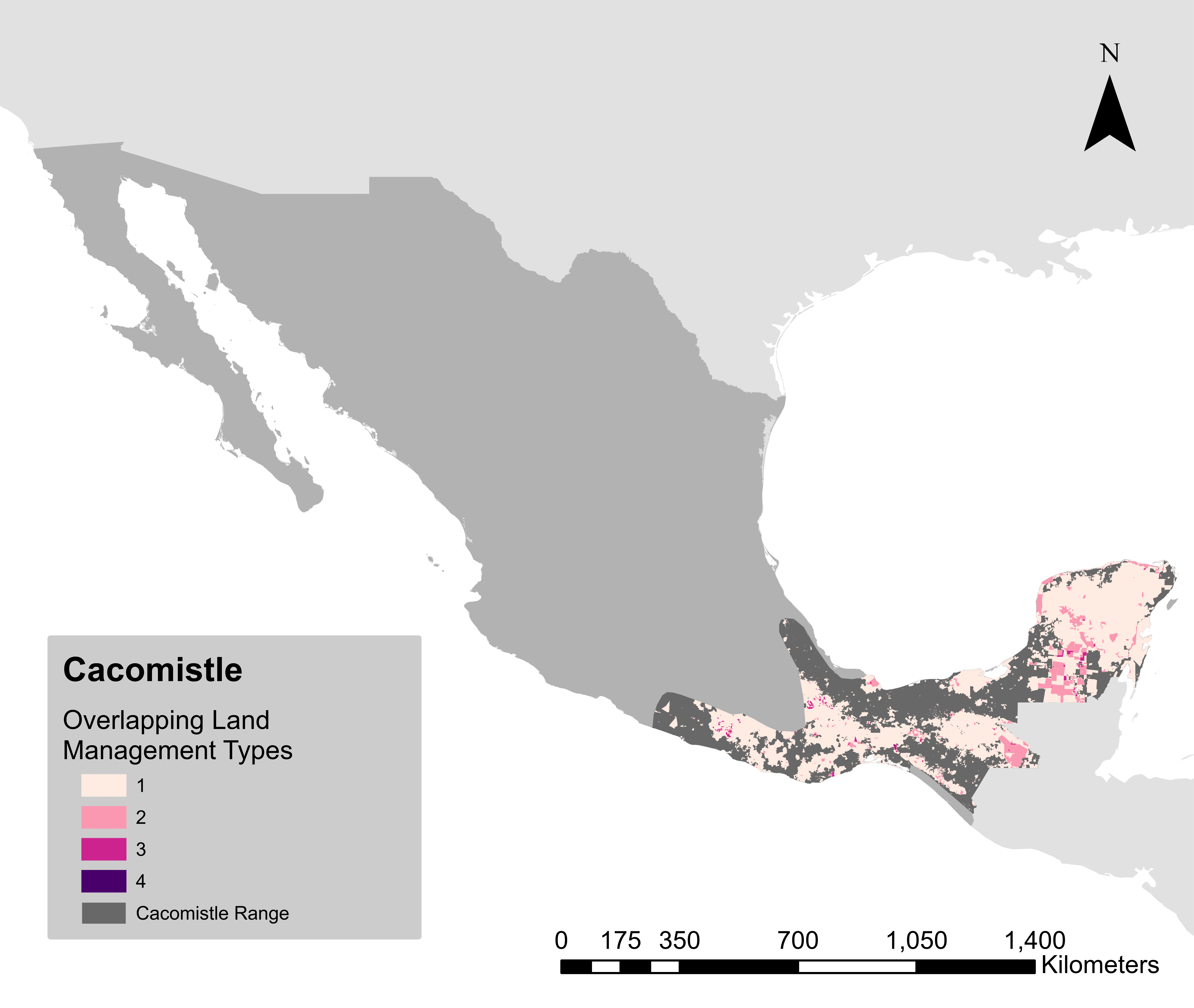

### CoyoteHM.png

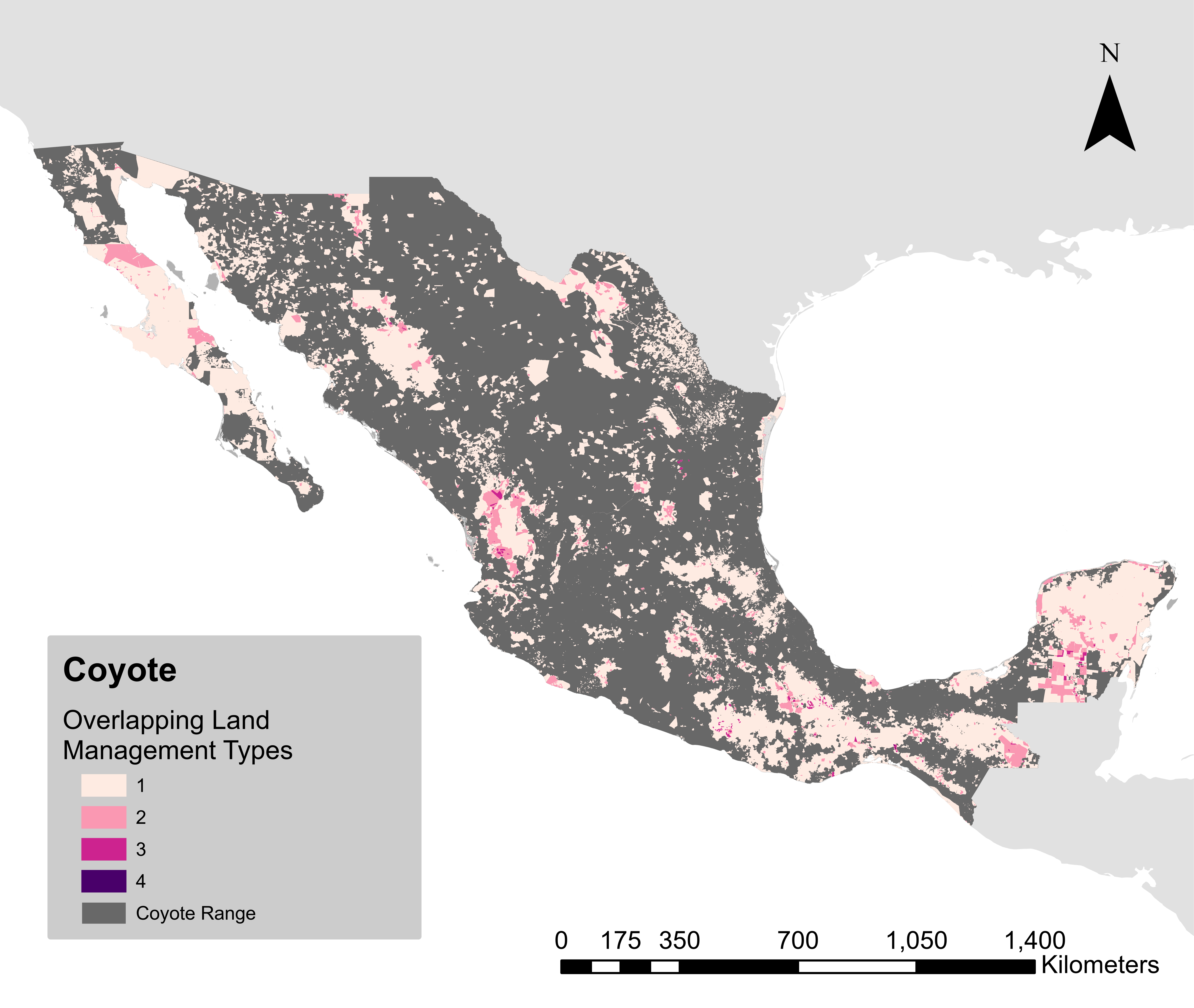

### EasternSpottedSkunkHM.png

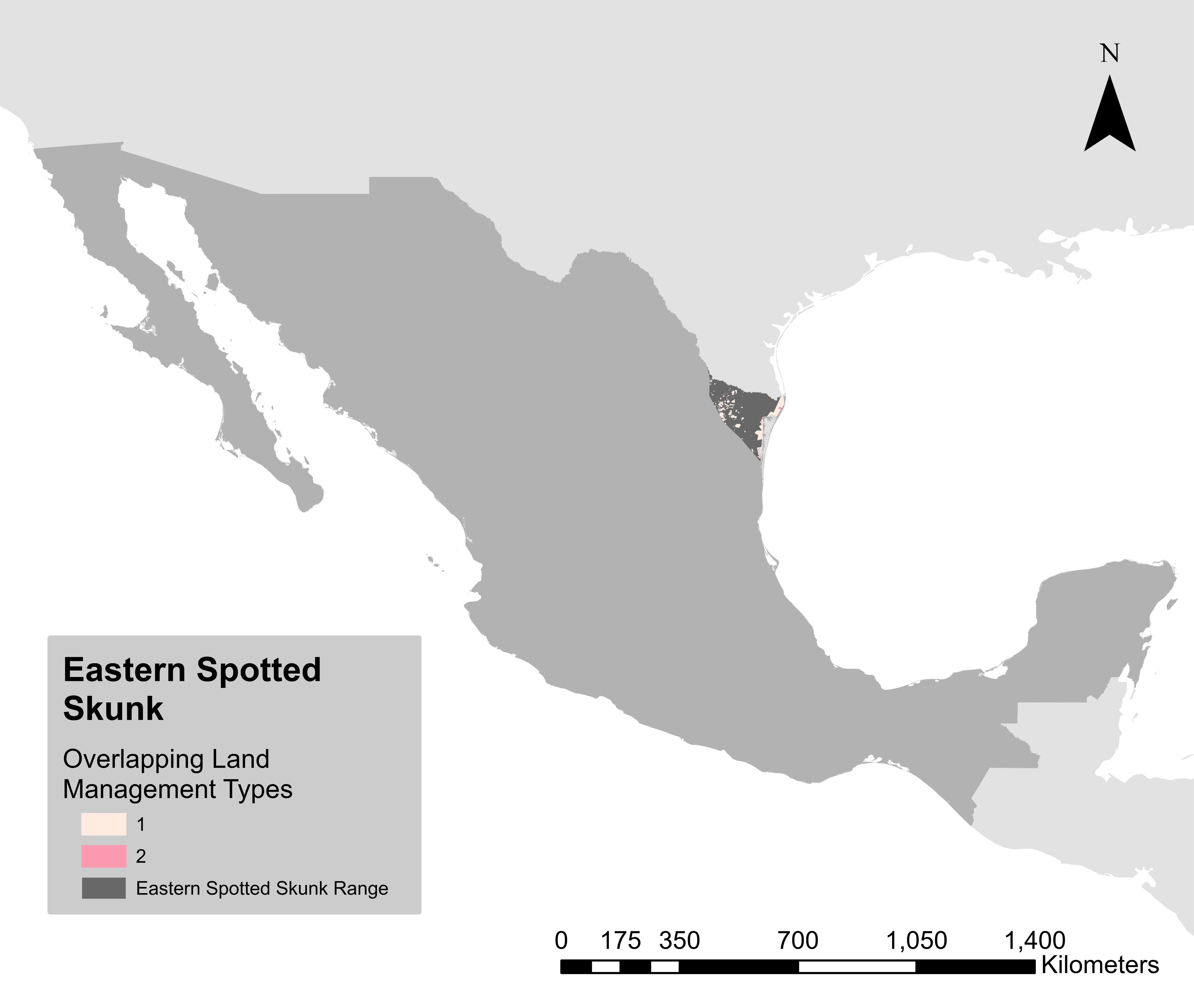

### GrayFoxHM.png

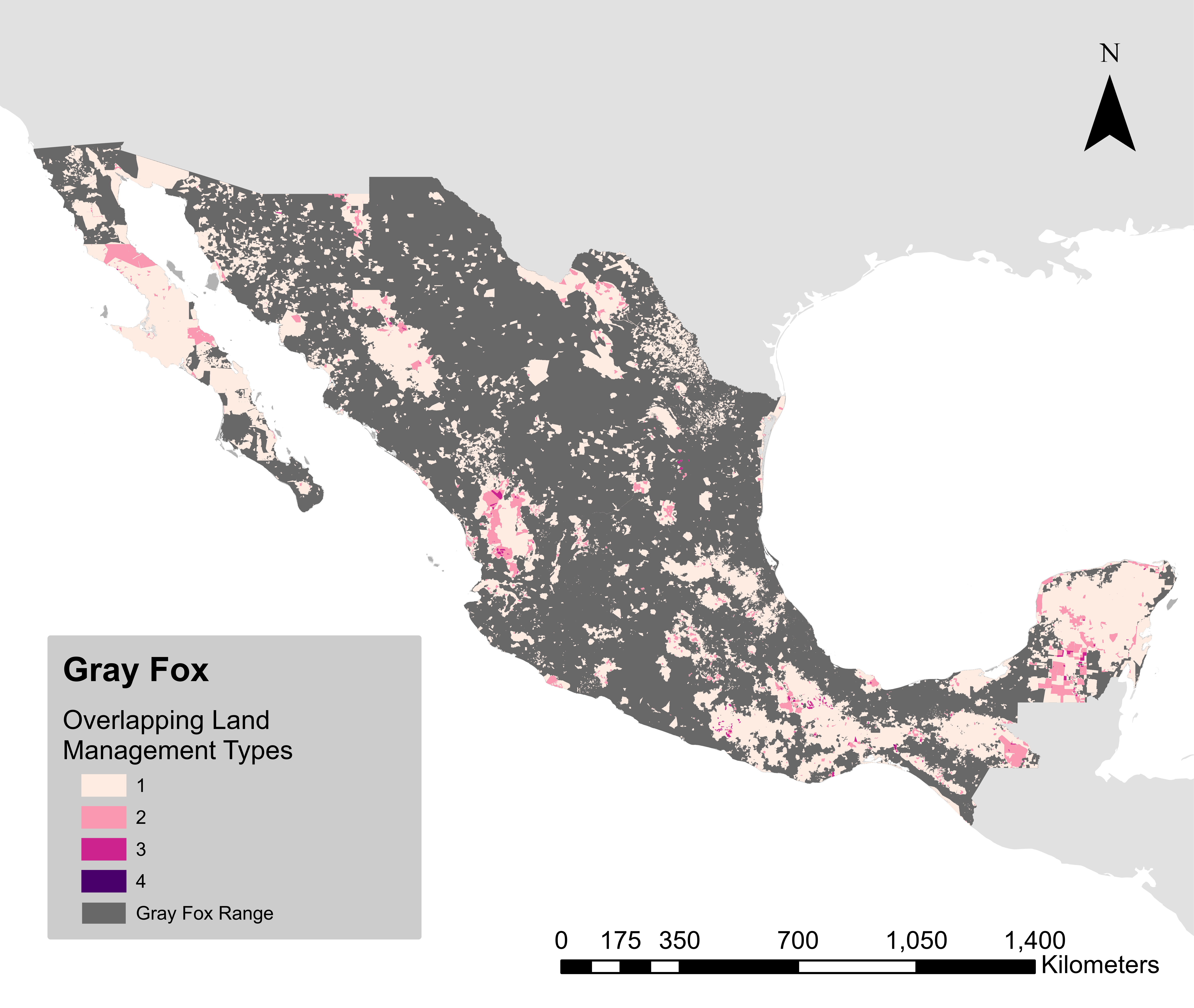

### GreaterGrisonHM.png

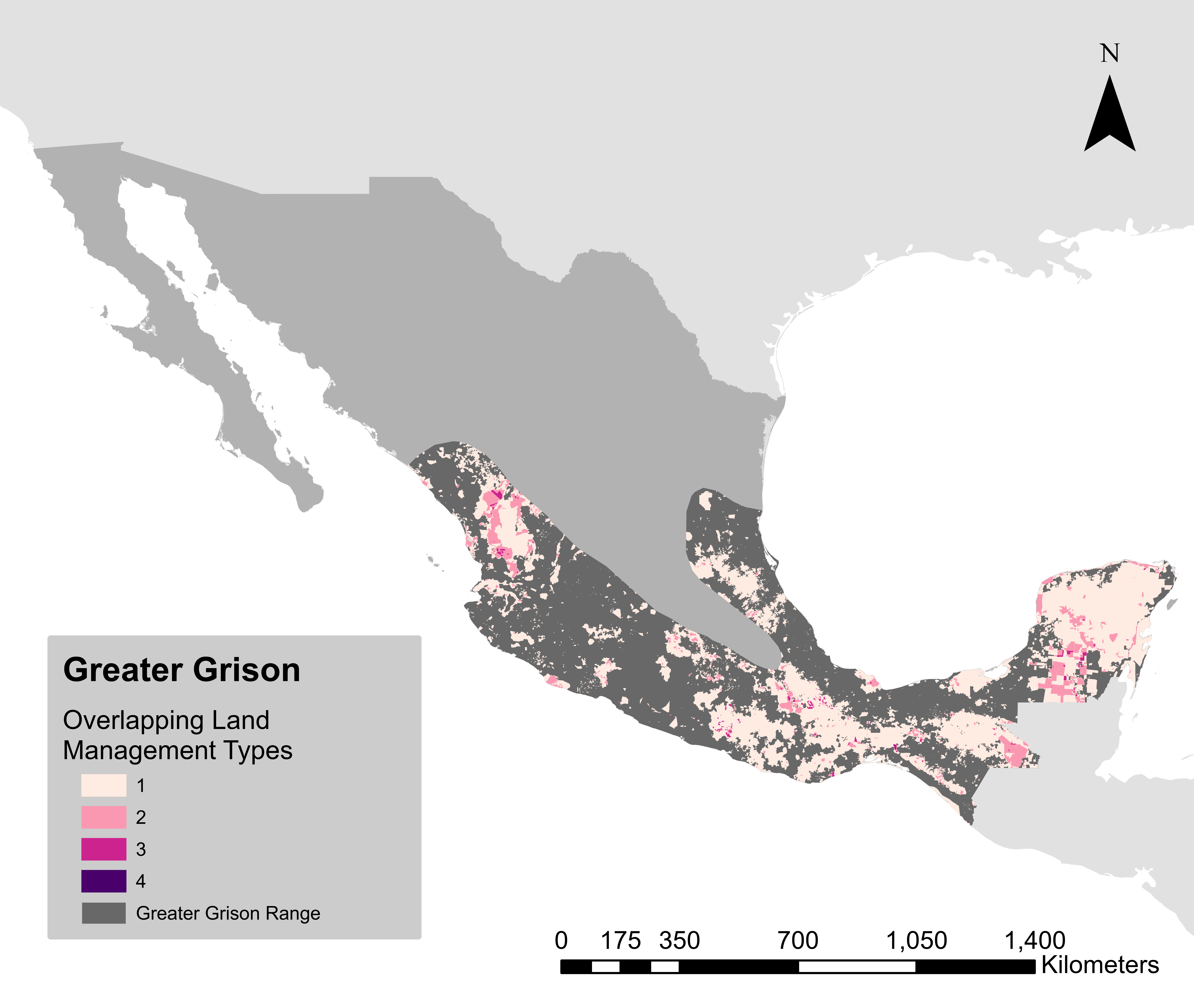

### HoodedSkunkHM.png

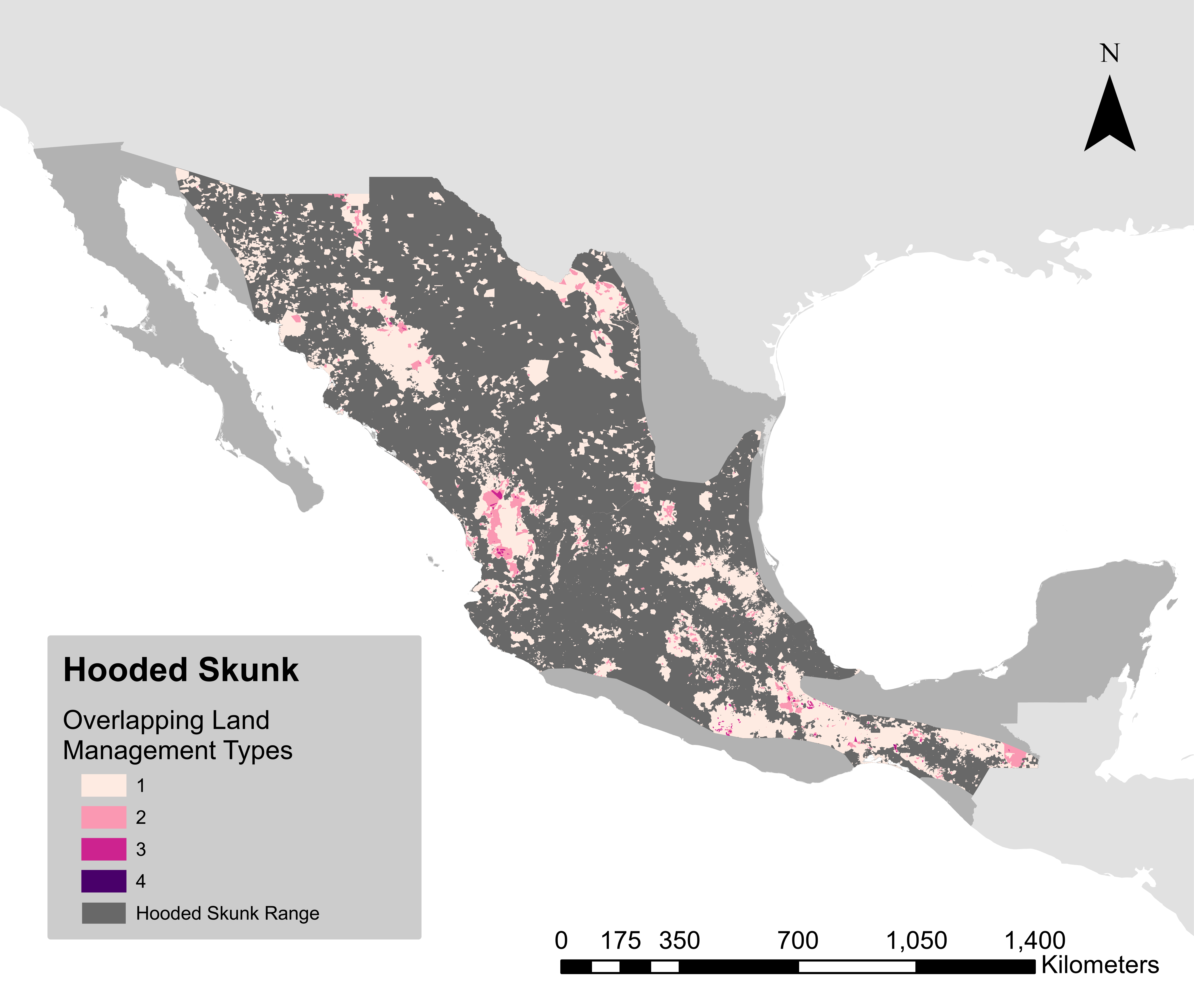

### JaguarHM.png

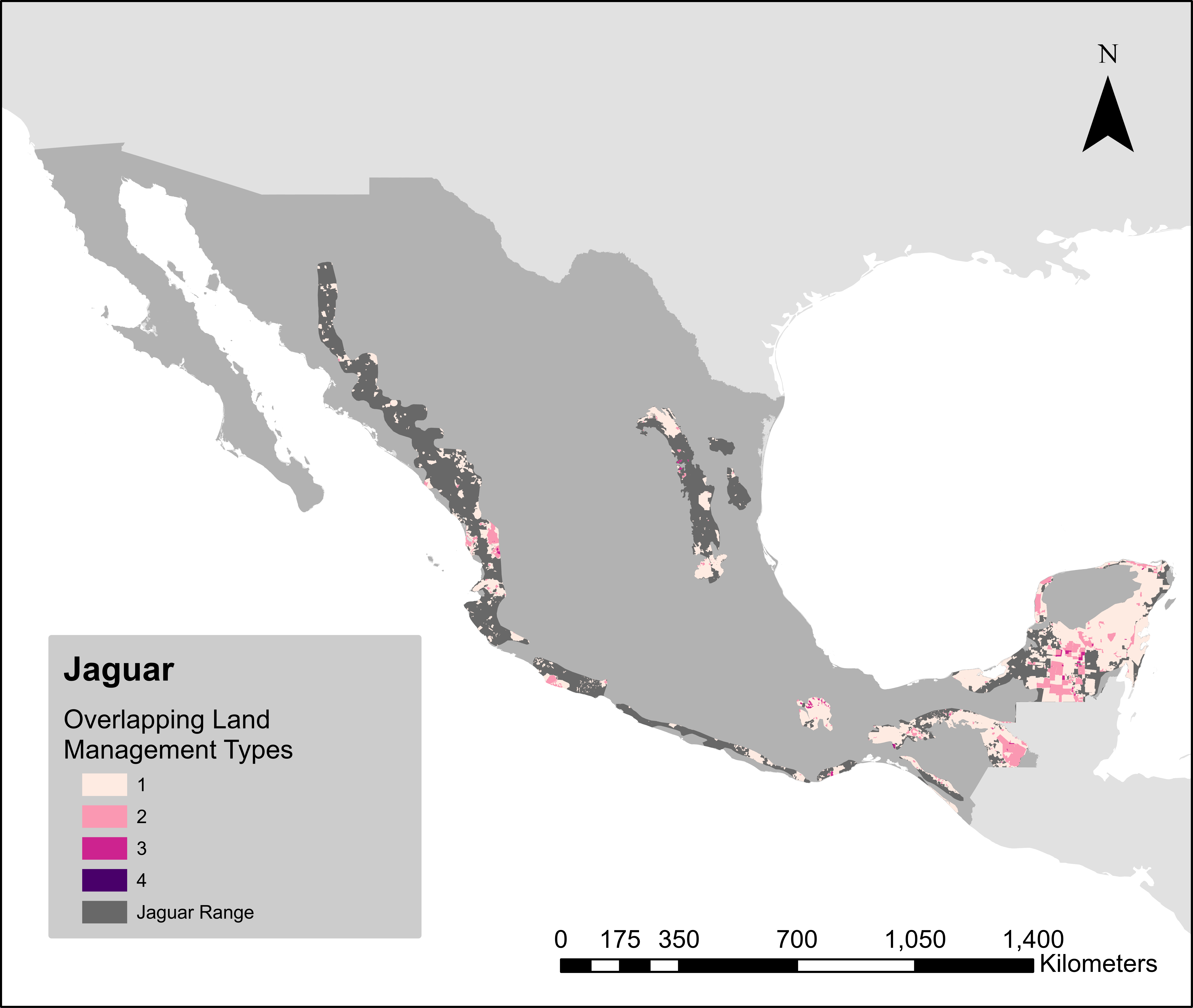

### JaguarundiHM.png

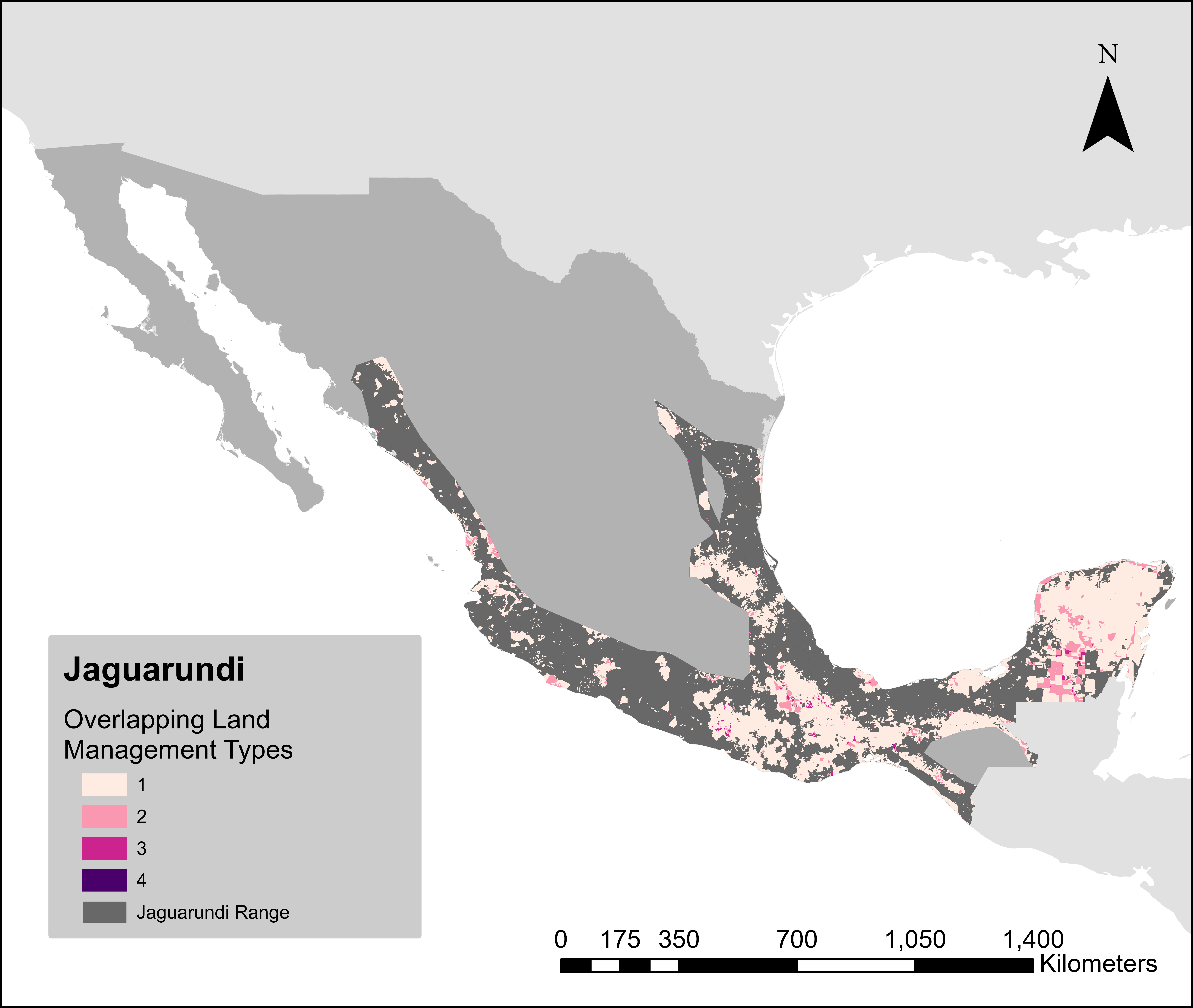

### KinkajouHM.png

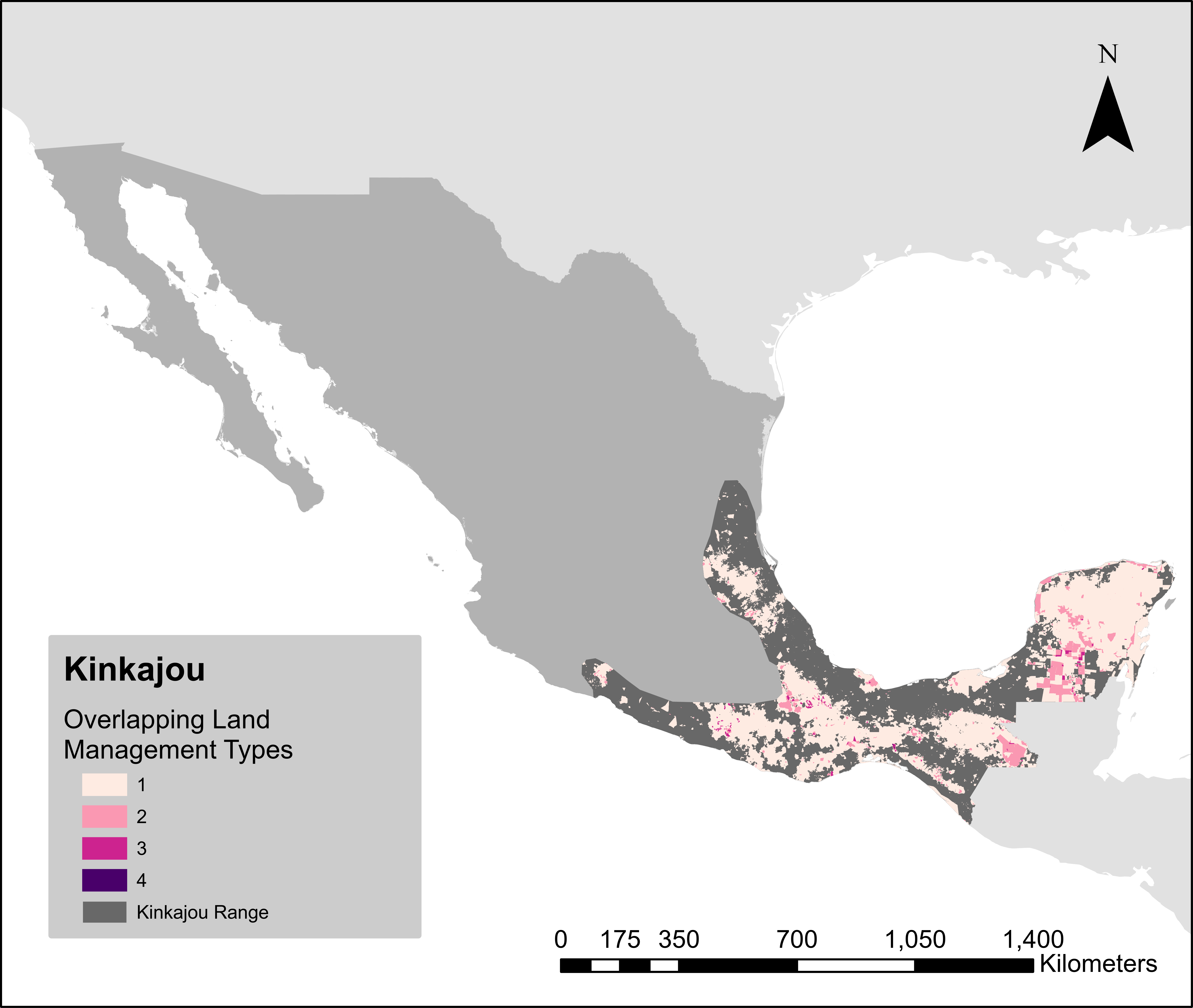

### KitFoxHM.png

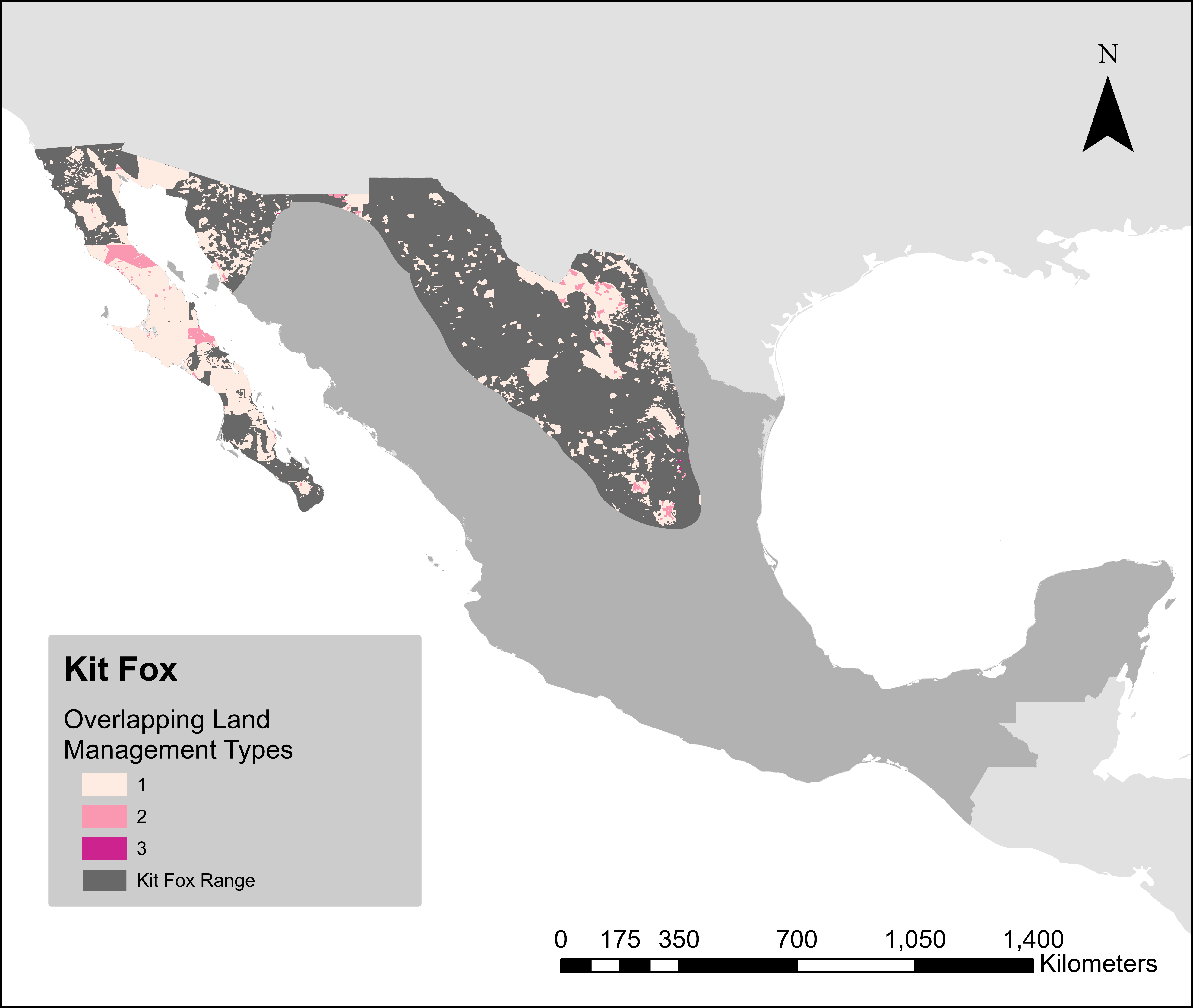

### LongTailedWeaselHM.png

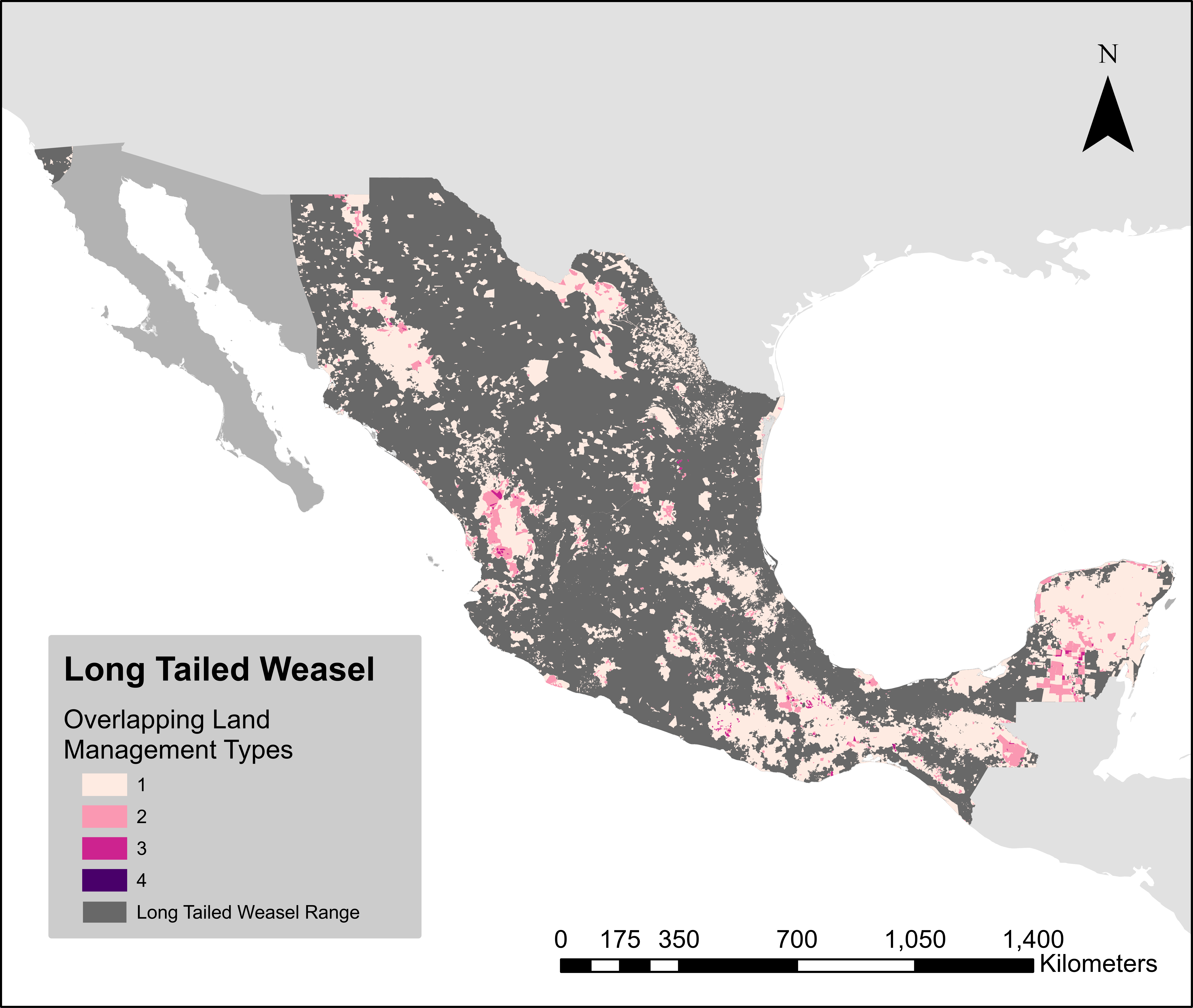

### MargayHM.png

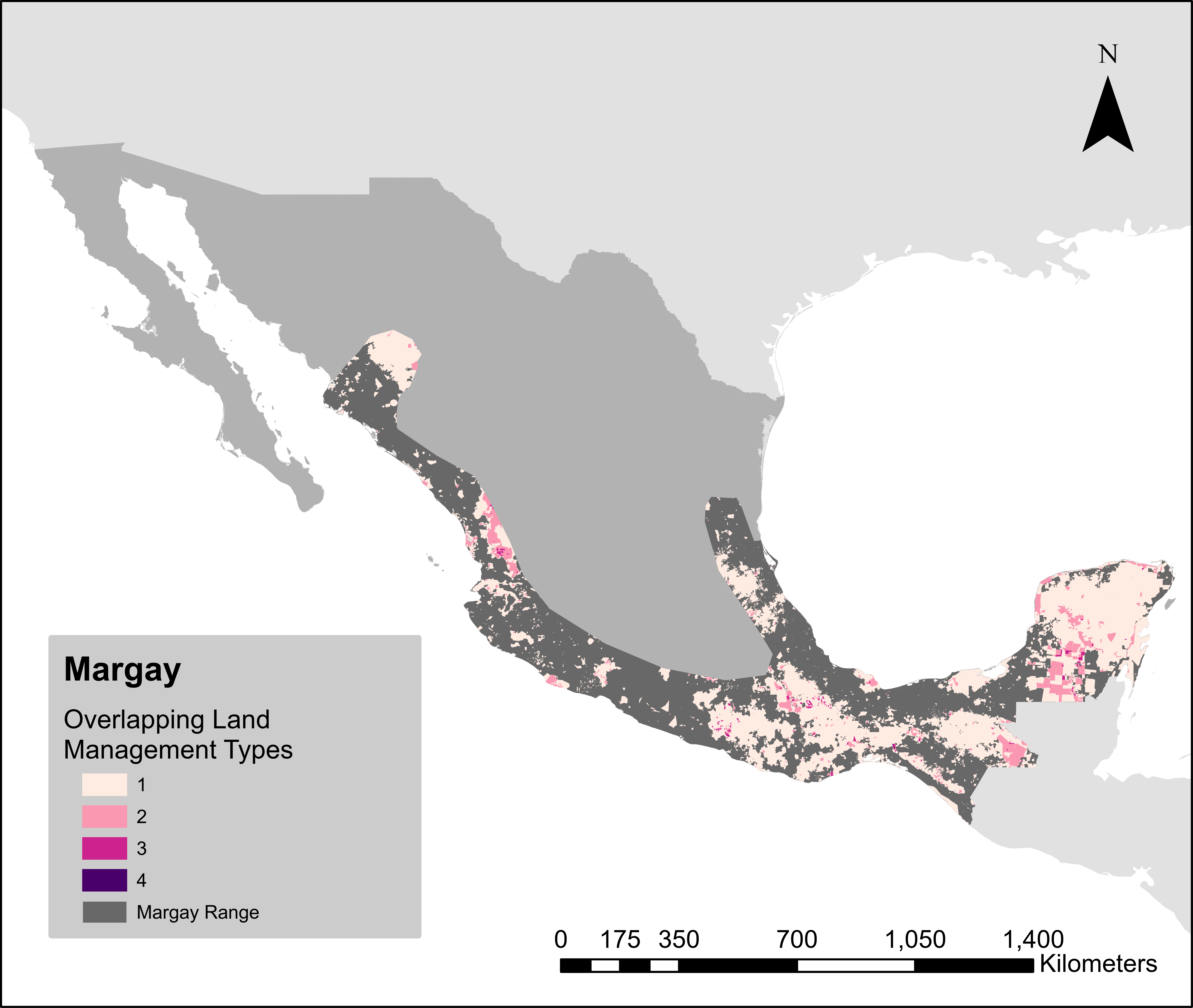

### OcelotHM.png

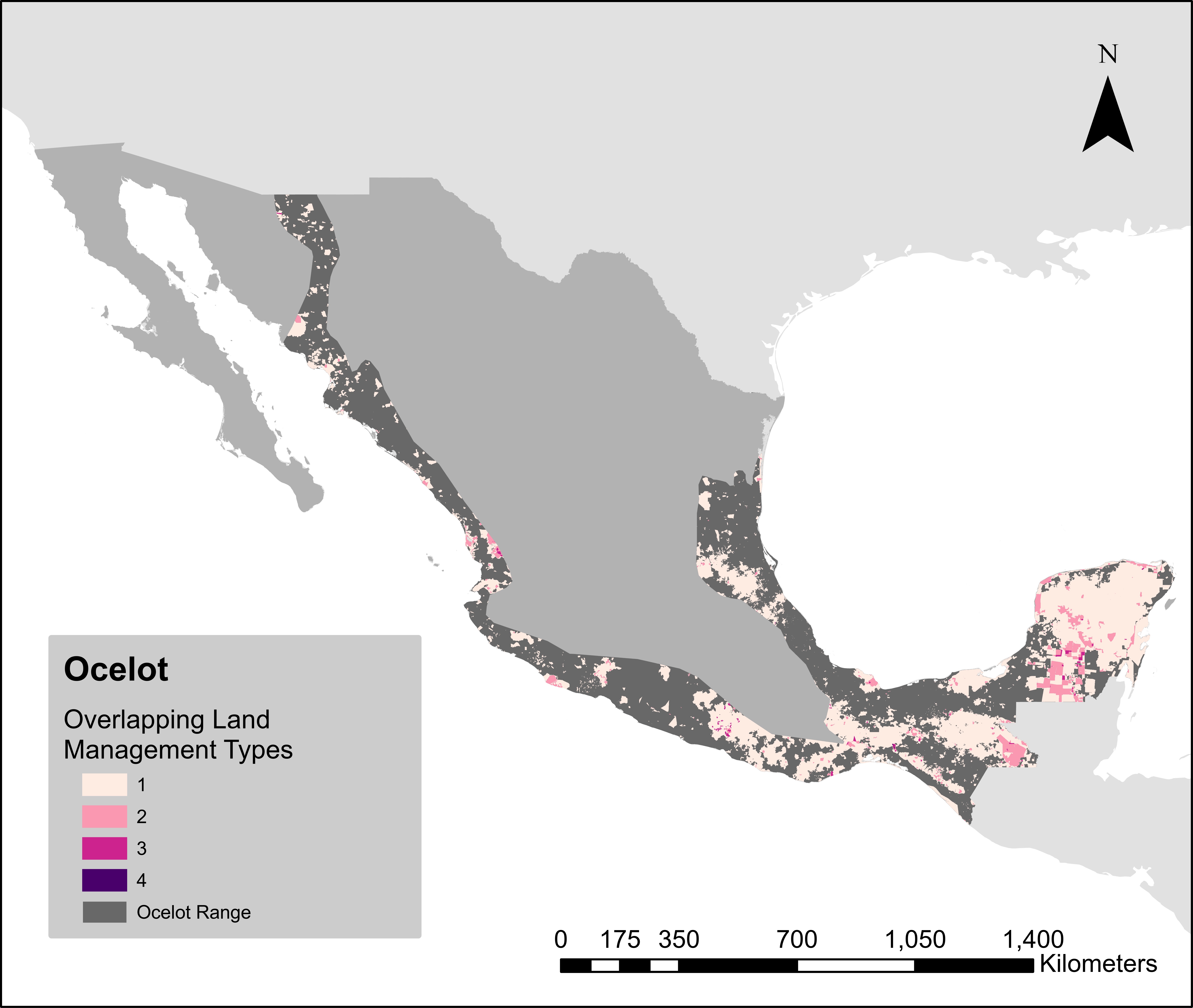

### PumaHM.png

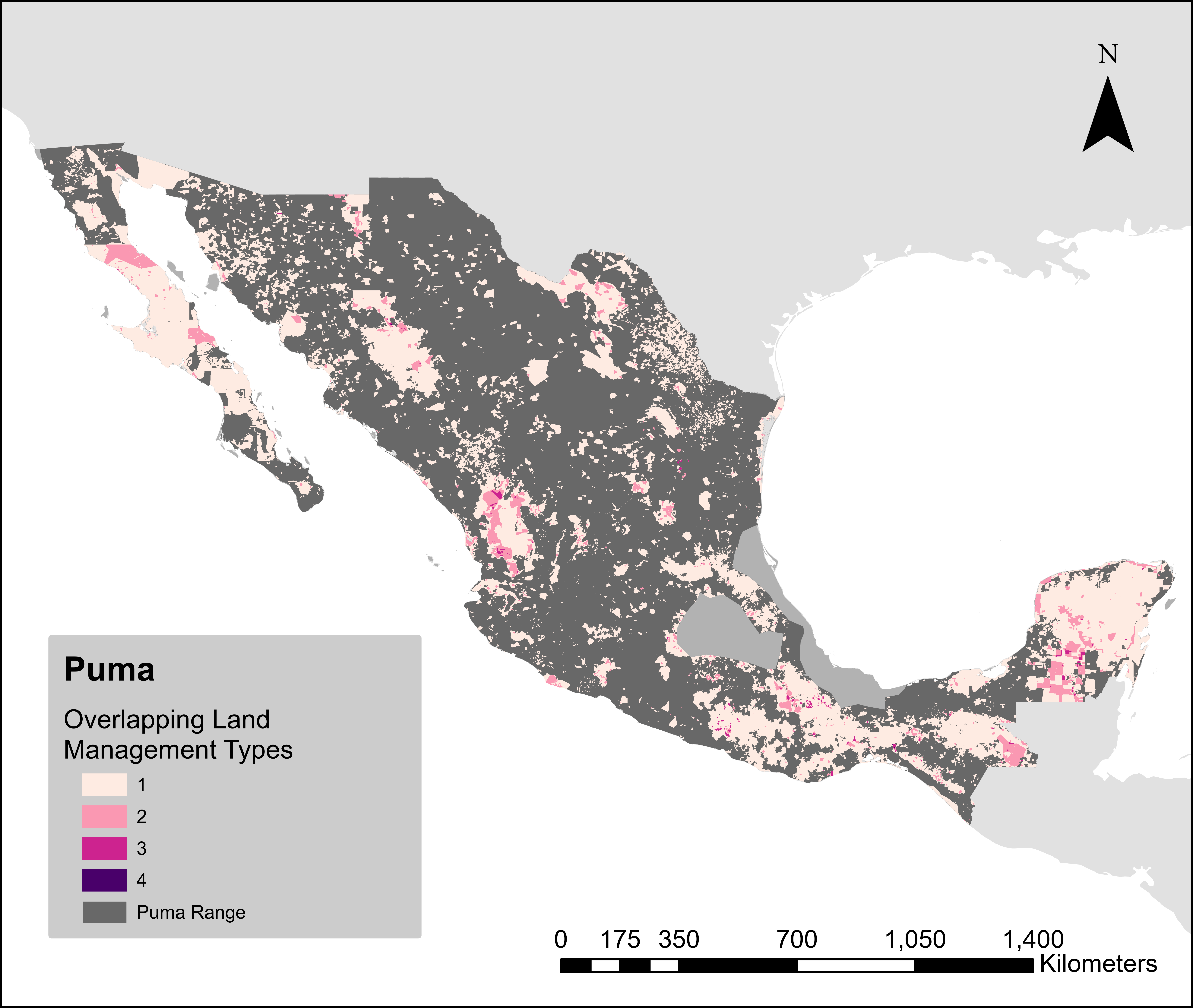

### PygmyRaccoonHM.png

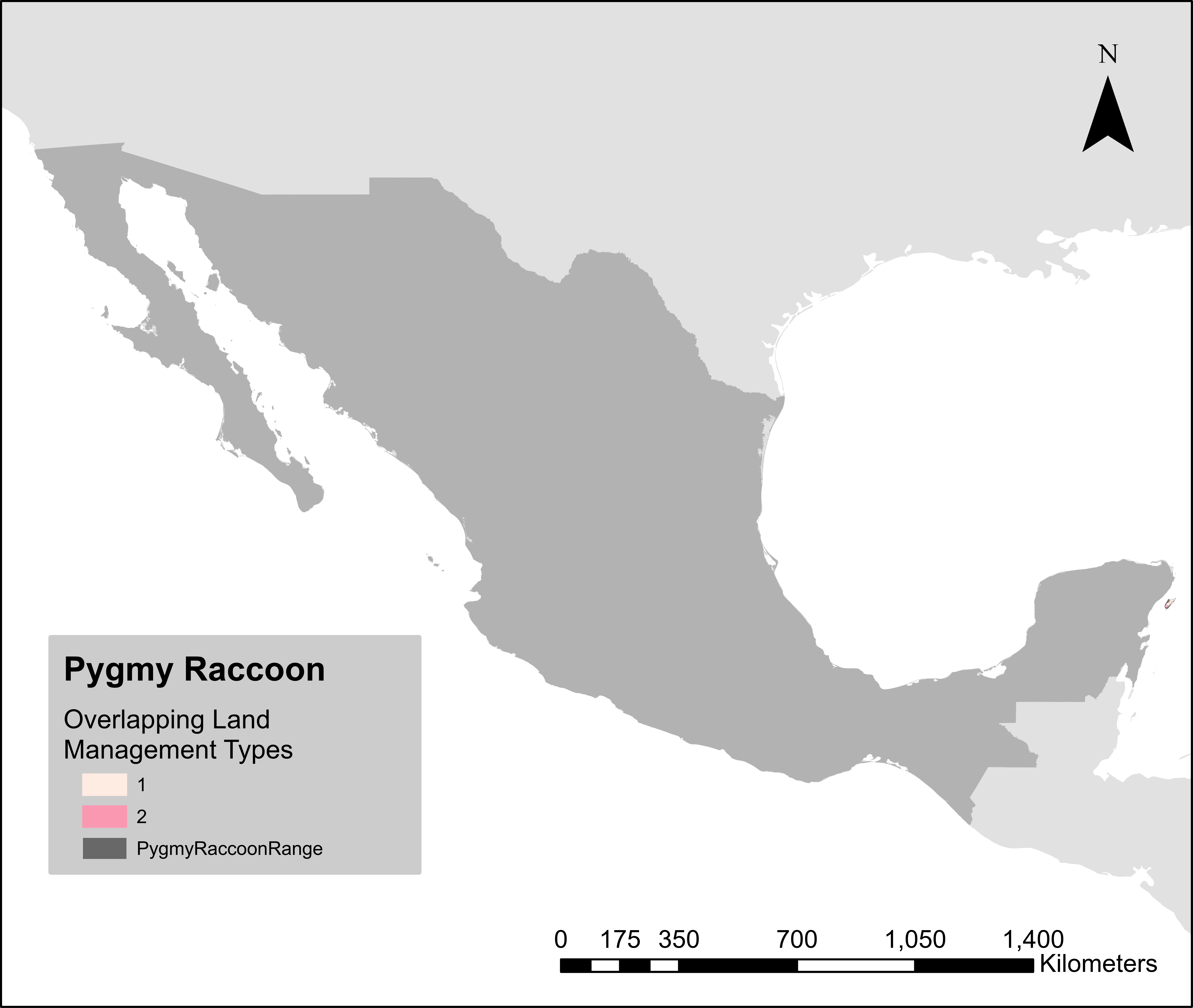

### PygmyRaccoonHM_Zoom.png

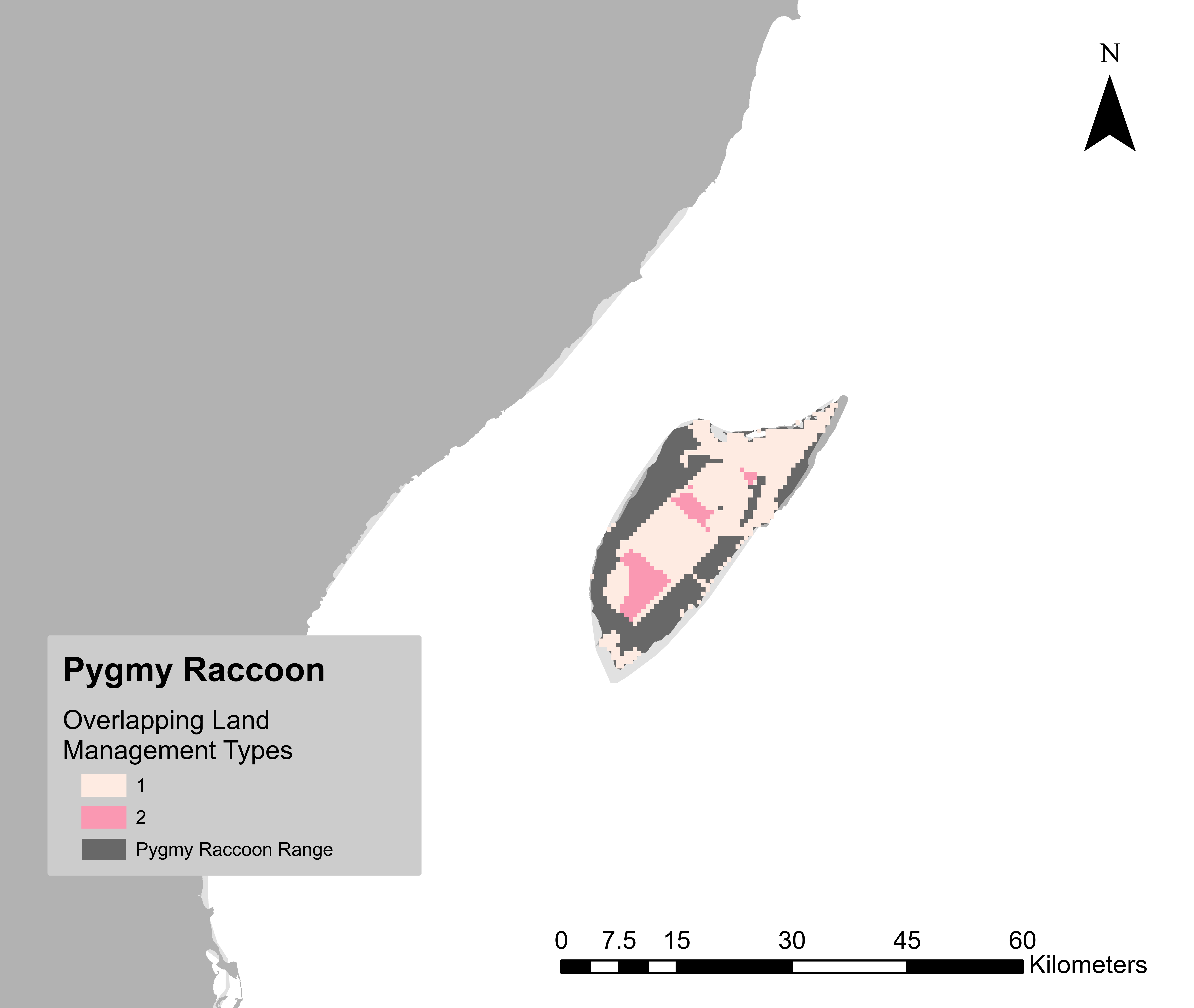

### PygmySpottedSkunkHM.png

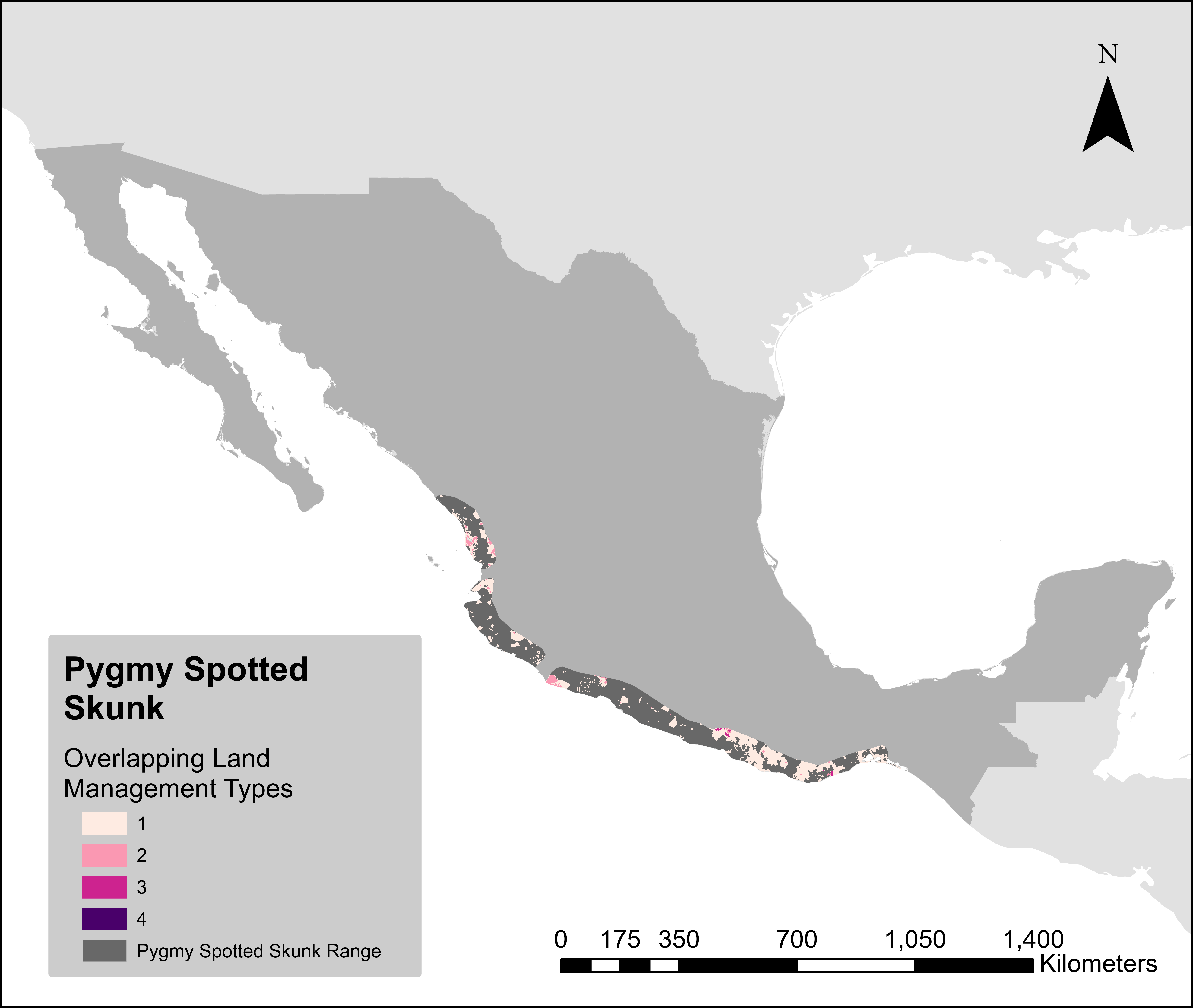

### RingtailHM.png

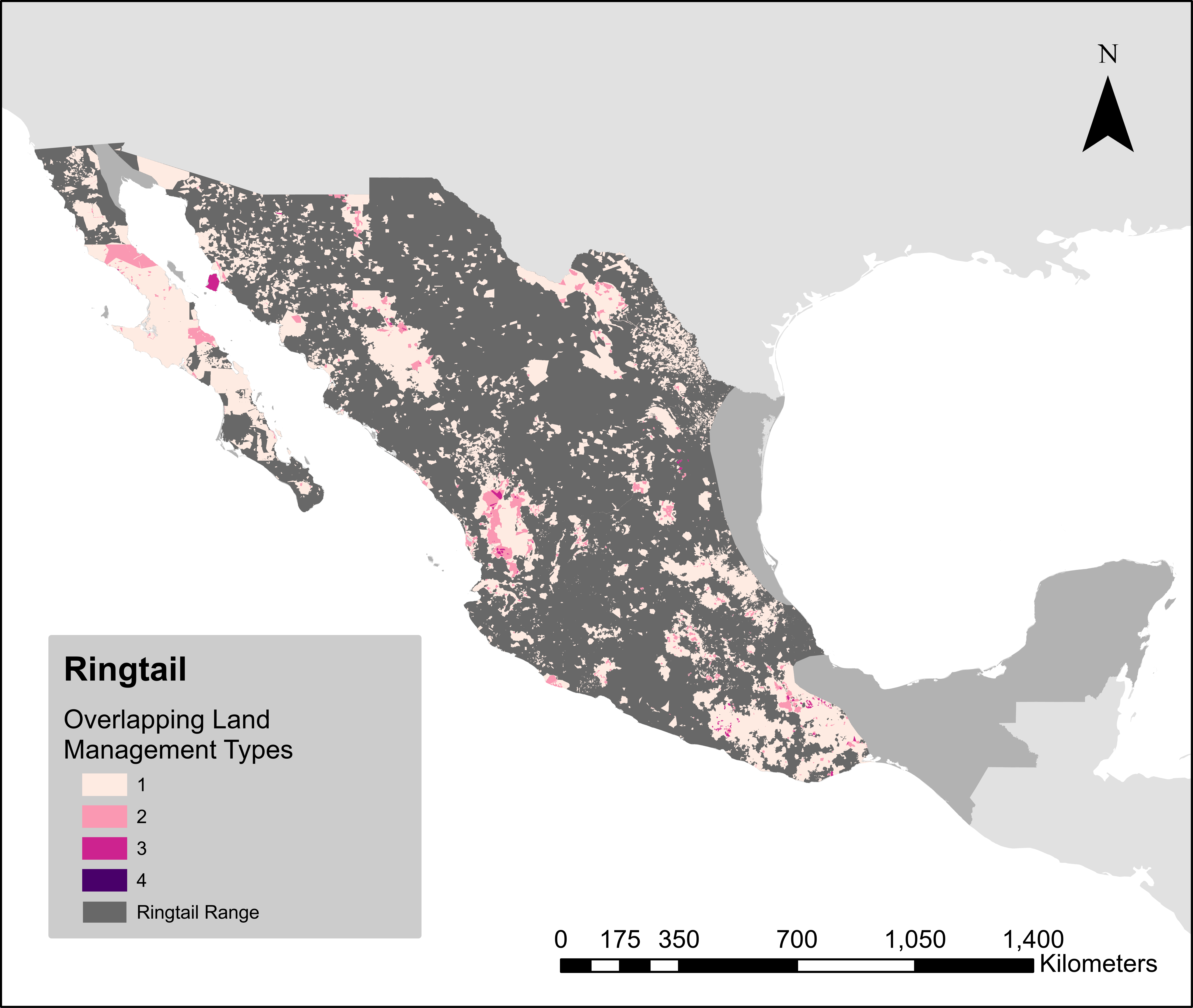

### SouthernSpottedSkunkHM.png

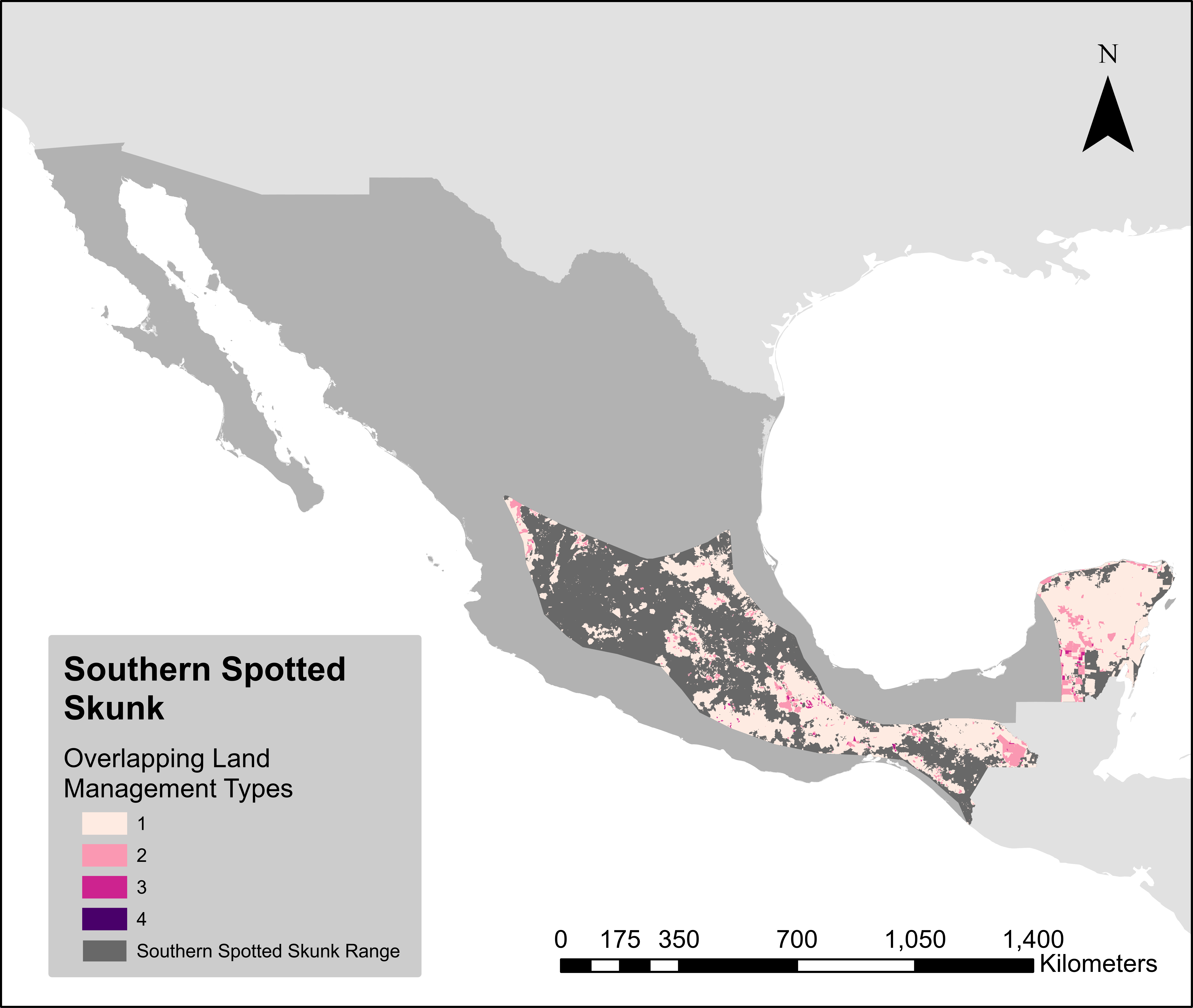

### StripedHogNosedSkunkHM.png

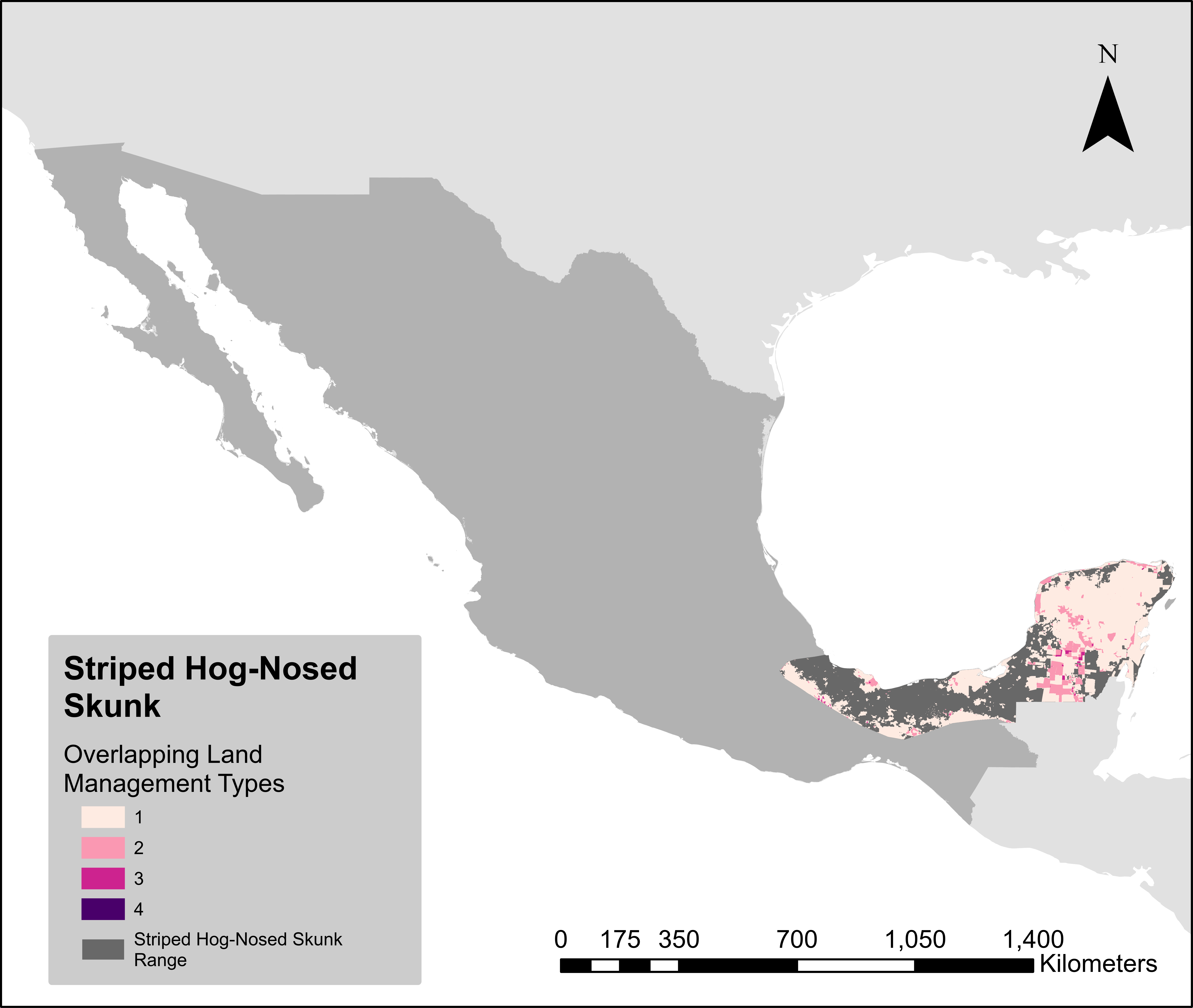

### StripedSkunkHM.png

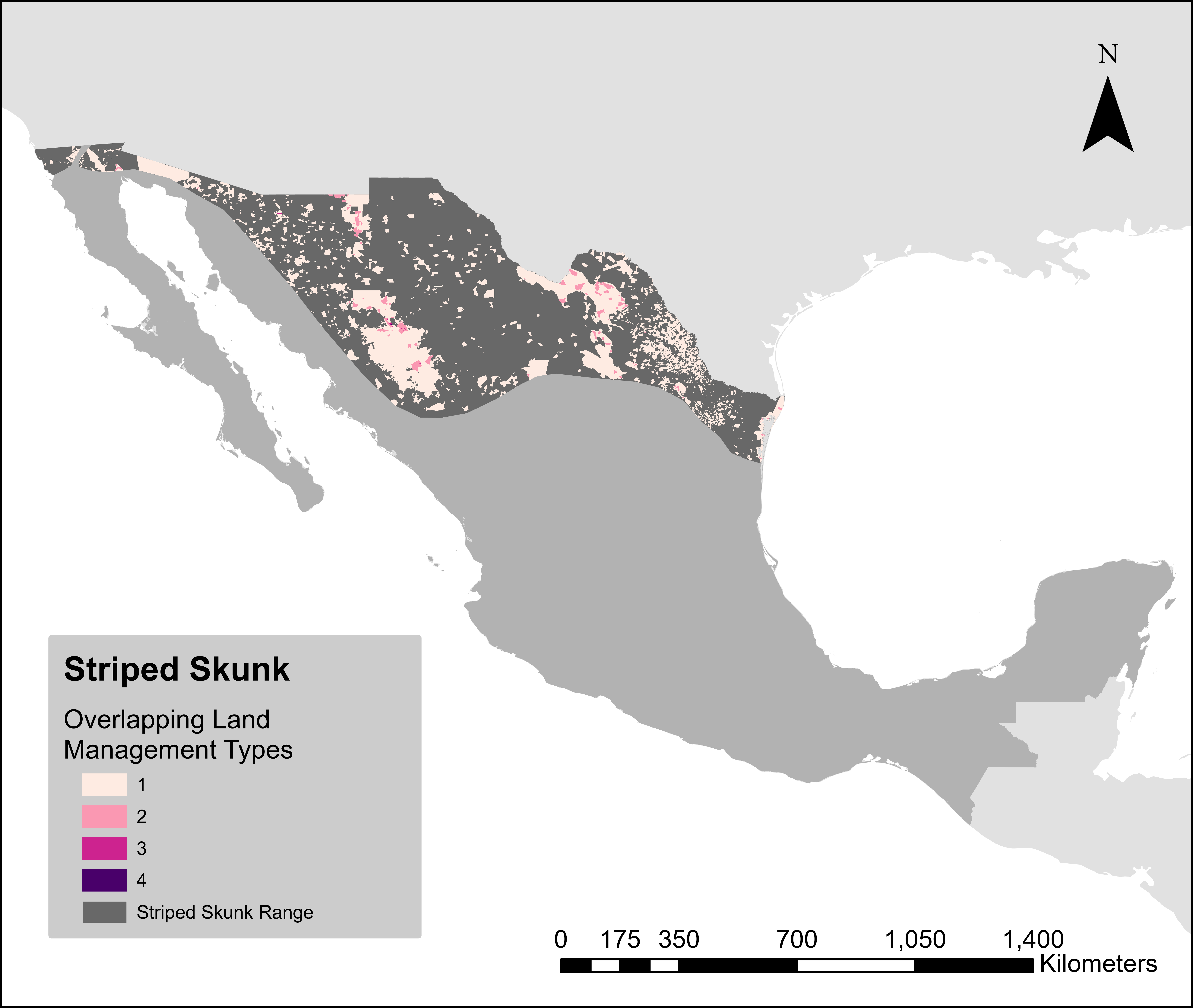

### TayraHM.png

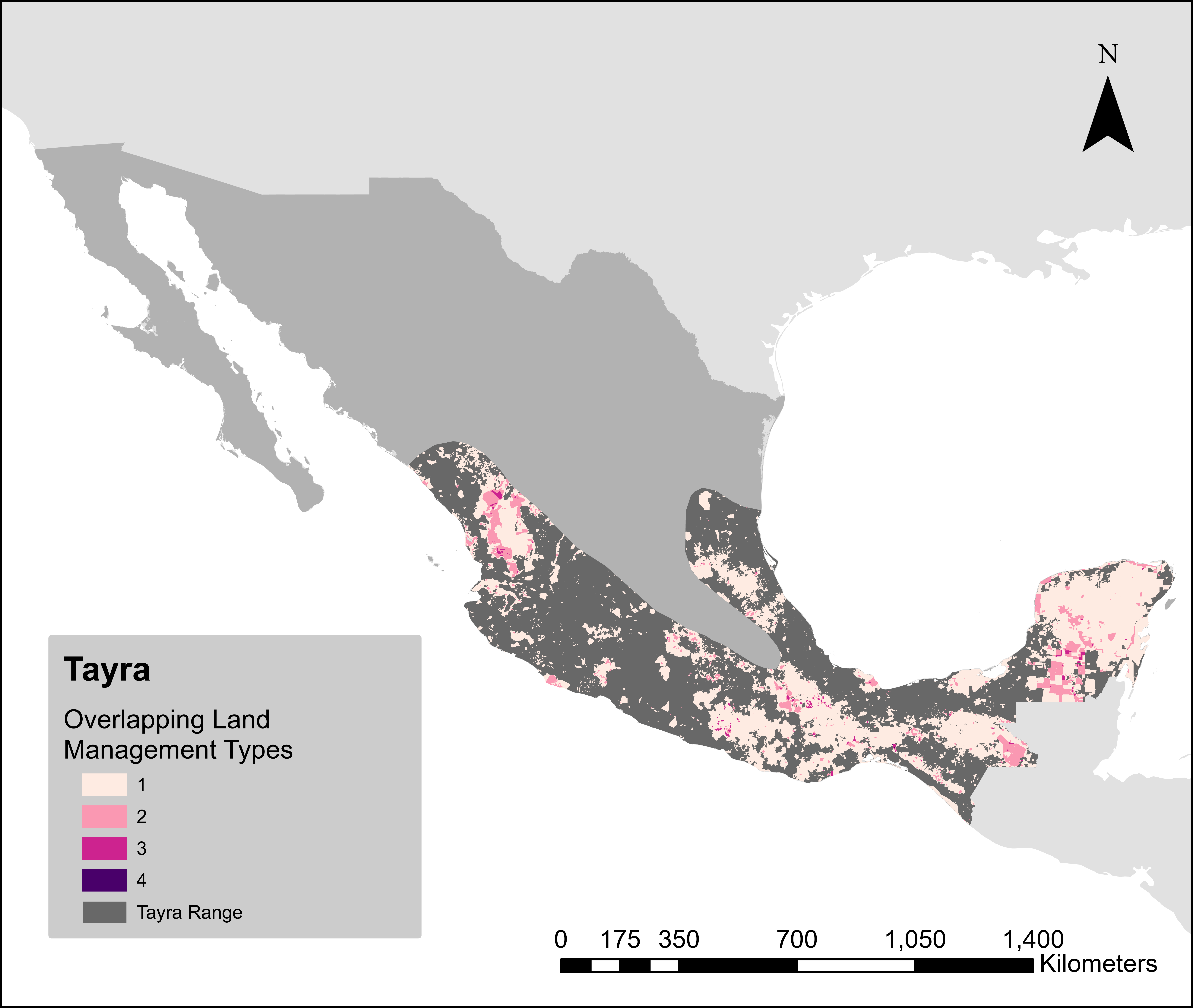

### WesternSpottedSkunkHM.png

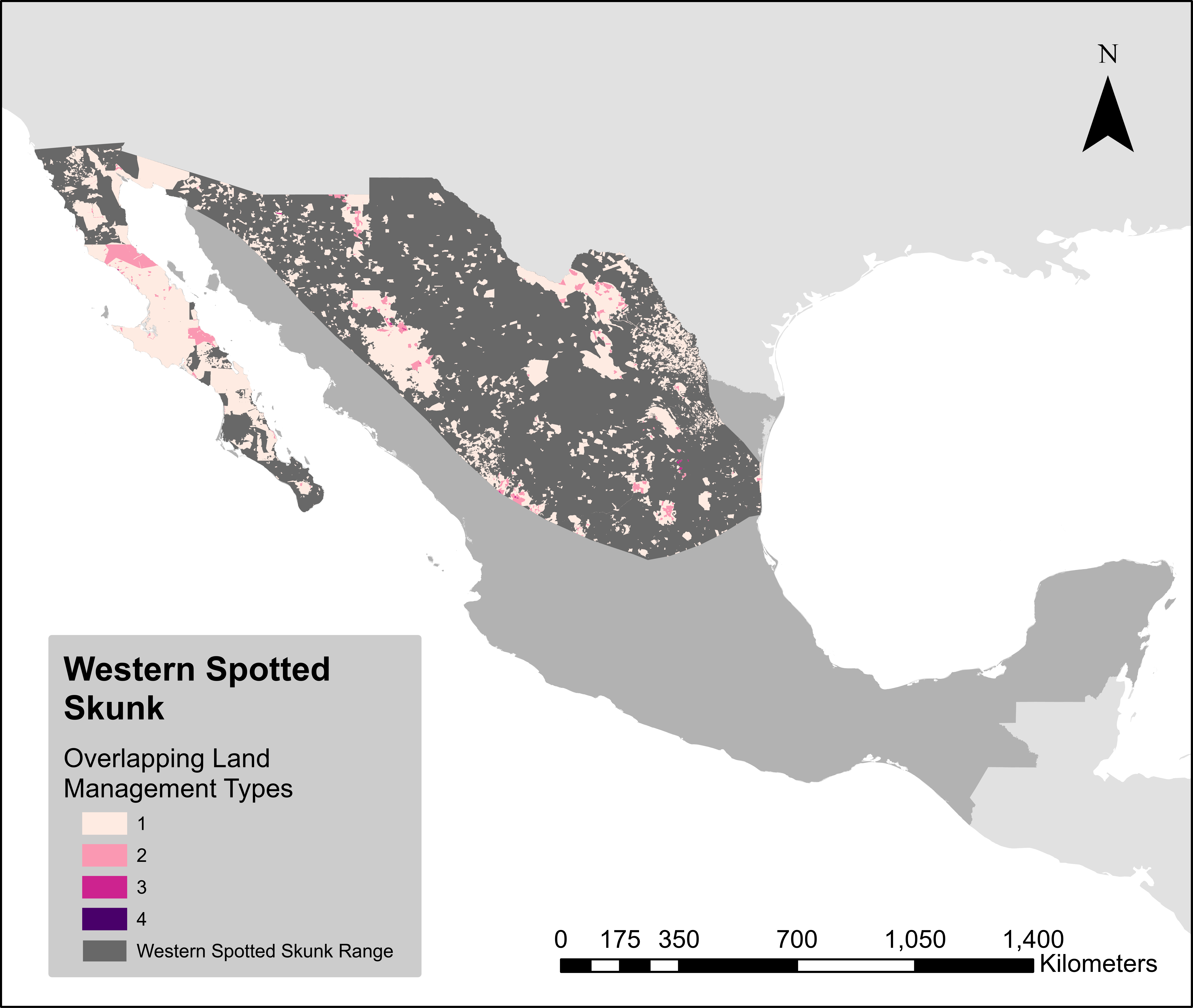

### WhiteNosedCoatiHM.png

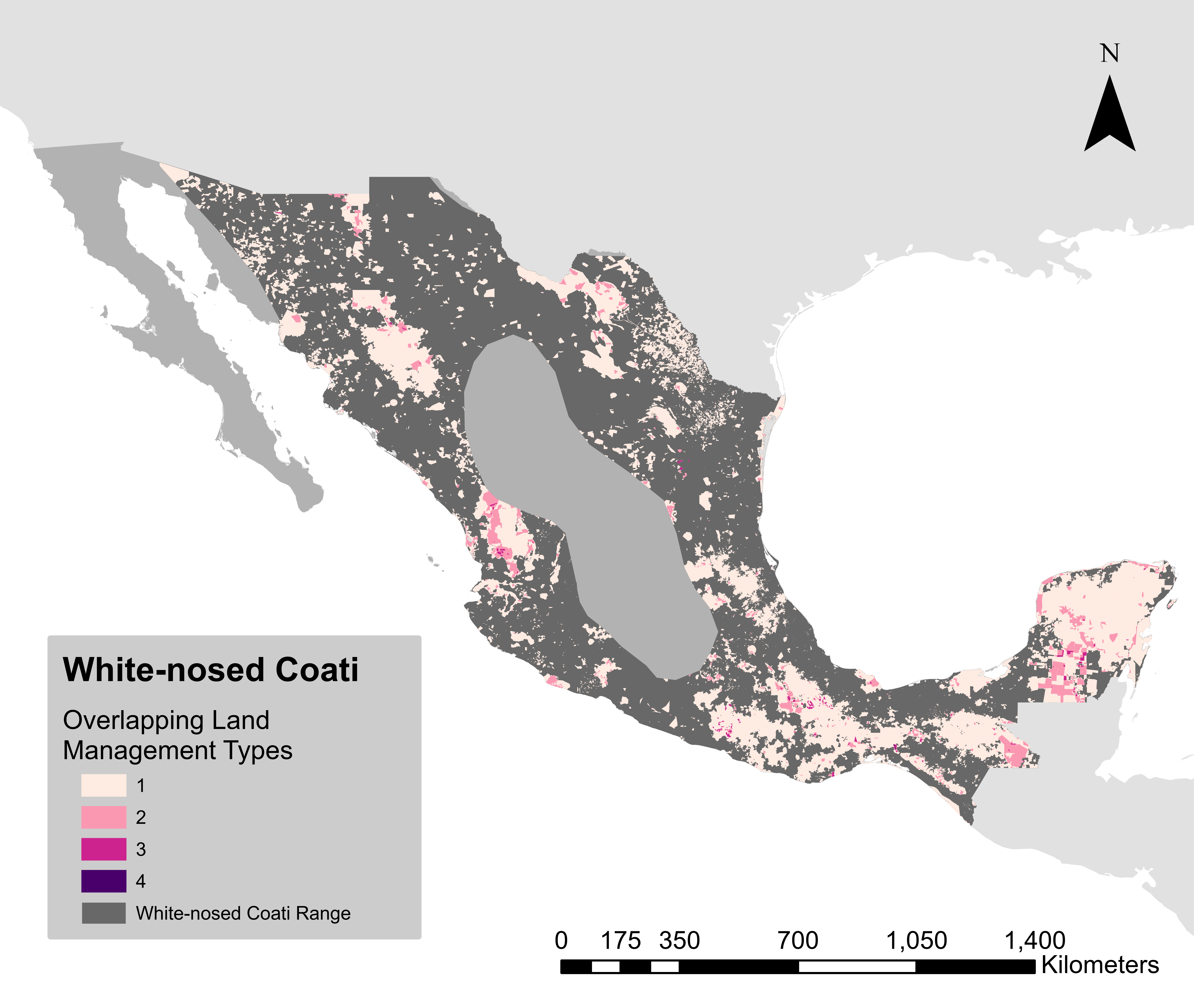
